# Effects of α-crystallin gene knockout on zebrafish lens development

**DOI:** 10.1101/2021.12.22.473921

**Authors:** Mason Posner, Kelly L. Murray, Brandon Andrew, Stuart Brdicka, Alexis Roberts, Kirstan Franklin, Adil Hussen, Taylor Kaye, Emmaline Kepp, Mathew S. McDonald, Tyler Snodgrass, Keith Zientek, Larry L. David

## Abstract

The α-crystallin small heat shock proteins contribute to the transparency and refractive properties of the vertebrate eye lens and prevent the protein aggregation that would otherwise produce lens cataracts, the leading cause of human blindness. There are conflicting data in the literature as to what role the α-crystallins may play in early lens development. In this study, we used CRISPR gene editing to produce zebrafish lines with null mutations for each of the three α-crystallin genes (*cryaa, cryaba* and *cryabb*). The absence of normal protein was confirmed by mass spectrometry, and lens phenotypes were assessed with differential interference contrast microscopy and histology. Loss of αA-crystallin produced a variety of lens defects with varying severity in larval lenses at 3 and 4 dpf but little substantial change in normal fiber cell denucleation. Loss of either αBa- or full-length αBb-crystallin produced no substantial lens defects. Mutation of each α-crystallin gene did not alter the mRNA levels of the remaining two, suggesting a lack of genetic compensation. These data confirm a developmental role for αA-crystallin in lens development, but the range of phenotype severity suggests that its loss simply increases the chance for defects and that the protein is not essential. Our finding that *cryaba* and *cryabb* mutants lack noticeable lens defects is congruent with insubstantial transcript levels in lens epithelial and fiber cells. Future experiments can explore the molecular consequences of *cryaa* mutation and causes of lens defects in this null mutant, as well as the roles of other genes in lens development and function.

## 1. Introduction

Alpha crystallins are small heat shock proteins that contribute to the high protein density that produces the refractive power of the vertebrate eye lens (Horwitz, 1992). Because of their ability to bind denaturing proteins and block aggregation, α-crystallins are thought to prevent lens cataracts, the leading cause of human blindness worldwide. Their expression in tissues outside the eye, especially the broadly expressed αB-crystallin, also suggests that they may play diverse physiological roles (Srinivasan et al., 1992). Numerous studies link αB-crystallin to the physiology and disease of muscular and nervous tissue (Bova et al., 1999).

Zebrafish have been a valuable model for investigating lens crystallin function. Zebrafish and human lenses are made of similar proteins, and the mechanisms underlying their development are conserved (Greiling et al., 2010; Posner et al., 2008). The production of large numbers of transparent embryos and the ease with which zebrafish can be genetically manipulated make them a good tool for examining crystallin function. The presence of two αB-crystallin paralog genes in zebrafish provides an opportunity to dissect the roles of this multifunctional single mammalian protein (Smith et al., 2006). The accessibility of early-stage embryos and fast development times make zebrafish ideal for studying the roles of α-crystallins in early lens development. These studies in zebrafish are more convenient and less costly than in mammalian systems such as mice.

Various genetic knockout and translation blocking techniques have been used to investigate the impacts of α-crystallin loss on zebrafish lens development. These experiments showed that loss of any of the three α-crystallin genes produced lens defects in 3- and 4-day-old fish (Mishra et al., 2018; Zou et al., 2015). This result differed from work in mice, which showed that while the loss of αA-crystallin led to reduced lens size and early cataract development, the loss of αB-crystallin did not produce a lens phenotype, although heart defects were found (Brady et al., 2001, 1997). Furthermore, our recent analysis of zebrafish gene expression at single-cell resolution indicated that neither of the two αB-crystallin genes were expressed in the lens at substantial levels through five days of development (Farnsworth et al., 2021). Therefore, there seem to be conflicting reports on what role, if any, αB-crystallins play in early lens development.

In the present study, we examined the larval lens in newly generated knockout lines for all three zebrafish α-crystallin genes. The resulting data address open questions about what effect, if any, the loss of either αB-crystallin protein will have on lens development. We also provide a more detailed analysis of lens phenotypes after αA-crystallin loss and compare several approaches for producing gene knockouts using CRISPR editing. This data analysis can provide a set of best practices for future studies of lens gene function.

## 2. Materials and Methods

### 2.1. Fish maintenance and treatment

All experiments performed with zebrafish in this study were approved by Ashland University’s Animal Use and Care Committee (approval #MP 2019-1) and the reporting in the manuscript follows the recommendations in the ARRIVE guidelines (Kilkenny et al., 2010). Methods of anesthesia and euthanasia, as described below, were consistent with the *Guidelines for Use of Zebrafish in the NIH Intramural Research Program* and were in accordance with the AVMA guidelines. ZDR strain zebrafish were maintained on a recirculating aquarium system at approximately 28°C with a 14:10 light and dark cycle with daily water changes. Juveniles and adults were fed twice each day with a combination of dry flake food and live *Artemia* cultures. Adults were bred in separate false bottom tanks to collect fertilized eggs, which were incubated in petri dishes with system water at 28°C. The resulting larvae that were to be raised to adulthood were fed a liquid slurry of ground zooplankton and green algae until large enough to eat flake food and *Artemia*.

### 2.2. Guide RNA design, production and injection

Target regions for the guide RNA (gRNA) used in single and double injections to produce null lines were identified using the CRISPR design tool in the *Benchling* digital notebook system (www.benchling.com). The target regions for the four-gRNA mix used to produce *cryaa* crispants were identified by Wu et al. 2018 (Wu et al., 2018). Sequences for all gRNA primers are shown in Supplementary Table S1. Guide RNAs were produced using a PCR-based plasmid-free approach that annealed and elongated a universal scaffold primer with one, two or four additional primers containing the sequence of the target region. Equimolar amounts of universal scaffold primer and target region primers were used with Q5 DNA polymerase (NEB). Cycling parameters included a touchdown PCR protocol: initial 94°C denaturation for 5 min; cycle parameters: 94°C for 20 sec, 65° for 20 sec, 68°C for 15 sec (13 additional cycles with extension temperature reduced 0.5°C per cycle); 30 additional cycles of 94°C for 20 sec, 58°C for 20 sec, 68°C for 15 sec; final extension 68°C for 5 minutes. The resulting PCR product was purified with the Monarch PCR and DNA Cleanup Kit (NEB) with elution in 20 microliters nuclease-free water. Up to 1000 ng of purified PCR product was used with the HighScribe T7 kit (NEB) and incubated overnight at 37°C, and the reaction was purified with the Monarch RNA Cleanup Kit (500 micrograms) with elution in 50 µl nuclease-free water (NEB). The quality of the gRNA prep was confirmed by agarose gel electrophoresis and quantified using a NanoDrop One (Thermo Fisher).

Each injection mix contained 250 ng/µl total of the purified gRNAs and 500 ng/µl of recombinant Cas9 protein (PNA Bio) with 0.2% phenol red in nuclease-free water. One nanoliter of mix was microinjected with a single 20 millisecond pulse into the cytoplasm of each zebrafish zygote using a Harvard Apparatus PL-90 picoinjector (Holliston, MA).

### 2.3. Identification of null lines

Embryos injected with single or double gRNAs were genotyped after at least two months of age. Fish were anesthetized in 0.016% tricaine methane sulfonate (MS-222) resuspended in trisma base, and the posterior edge of the fin was removed with scissors. Fish were kept separated for recovery and during genotyping. Genomic DNA was collected from each fin clip using the Monarch Genomic DNA Purification kit (New England Biolabs) and amplified using OneTaq 2X Master Mix (NEB) with PCR primers that flanked the gRNA-targeted region. The recommended PCR protocol was followed, with appropriate annealing temperatures determined experimentally. All primers and annealing temperatures are shown in Supplementary Table S1. Amplified genomic DNA targeted with one gRNA was sequenced to determine whether more than one sequence was apparent beginning at the target area. Mutated genomic regions targeted with two gRNAs could be identified directly by the size of the resulting PCR product and then sequenced to confirm the resulting allele.

Fish resulting from injected zygotes (founder fish: F0) determined through fin clipping to contain mutant alleles were outcrossed to wild-type fish. Ten resulting embryos were combined and homogenized to purify genomic DNA, and the targeted gene region was amplified to determine if the parent founder fish had passed on mutant alleles to offspring. If offspring were found to be heterozygous for a gene mutation, larvae were raised another two months for fin clipping. For single gRNA-injected fish, sequencing of the heterozygotes was performed to determine if the resulting mutation would cause a frame shift and early stop codon. Multiple heterozygotes with the same identified frameshift mutation were incrossed to produce null individuals that were then used to produce null lines. For double gRNA-injected embryos, sequencing was used to identify heterozygotes with the same deletion, which were then incrossed to generate a null line.

### 2.4. Proteomic assessment of knockout fish

A detailed protocol outlining the sample preparation of zebrafish lenses to detect the presence of α-crystallins was previously published (Posner et al., 2017). Briefly, 50 µg portions of protein derived from either one or two dissected lenses were digested overnight with trypsin in the presence of ProteaseMax™ detergent as recommended by the manufacturer (ProMega, Madison, WI, USA). One microgram of each tryptic digest was then injected at 10 µl/min onto a Symmetry C18 peptide trap (Waters Corporation, Milford, MA, USA) for 0-5 min using 98% water, 2% acetonitrile (ACN), and 0.1% formic acid mobile phase and then switched onto a 75 µm x 250 mm NanoAcquity BEH 130 C18 column (Waters Corporation) using mobile phases water (A) and ACN (B), each containing 0.1% formic acid.

Peptides were separated using 7.5-30% ACN over 5-35 min, 30-98% from 35-36 min, 98% ACN from 36-41 min, 98-2% ACN from 41-42 min, and 2% ACN from 41-60 min at a 300 nL/min flow rate. Eluent was analyzed using a Q-Exactive HF mass spectrometer using electrospray ionization with a Nano Flex Ion Spray Source fitted with a 20 µm stainless steel nanobore emitter spray tip and 1.0 kV source voltage (Thermo Fisher Scientific, San Jose, CA, USA). Xcalibur version 4.0 was used to control the system. Prior to the targeted parallel reaction monitoring data collection, a digest from an adult 12-month old wild-type lens was analyzed by data-dependent acquisition (DDA) to produce a zebrafish spectral library to design a targeted parallel reaction monitoring (PRM) method to measure the abundance of αA, αBa, and αBb-crystallins in wild-type and mutant fish. Precursor mass spectra were acquired over m/z 375 to 1,400 m/z at 120,000 resolution, automatic gain control (AGC) target 3 × 10^6^, maximum ion time (MIT) of 50 ms, profile mode, and lock mass correction using m/z = 445.12002 and 391.28429 polysiloxane ions. MS2 scans were acquired from 200-2000 m/z, intensity threshold of 5 × 10^4^, automatic gain control (AGC) 1 × 10^5^, maximum ion time (MIT) 100 ms, 30,000 resolution, profile mode, normalized collision energy (NCE) 30, isolation window of 1.2 m/z, loop count 10, and exclusion of +1 and unassigned charges. MS2 results were searched using Sequest within Protein Discoverer 1.4 (Thermo Fisher Scientific) using a tryptic search, 10 ppm precursor mass tolerance, 0.1 Da fragment ion tolerance, dynamic modification for oxidation of methionine, and a static modification of cysteines for carbamidomethyl alkylation. A uniprot filtered proteome 000000437 zebrafish protein database containing 43,607 entries downloaded on 5/17/2022 was used. Control of peptide false discovery used a reversed sequence strategy and Percolator software to calculate q scores. Skyline (v.22.2.0.255) (Pino et al., 2020) was then used to create a Biblio Spec spectral library containing 564 peptide entries using a cutoff score of 0.95. These peptides were associated with the above uniprot zebrafish database to match to 434 proteins and peptides matching αA-, αBa-, and αBb-crystallins were added to the target list. The raw file used to create this spectral library was then imported into Skyline and 3 peptides from each protein were selected that had both strong precursor intensities and complete fragment ion series. This precursor list was exported to create a PRM method within the instrument control software with identical chromatographic conditions as used in the DDA run. Precursor scans from 400-1000 m/z at 120,000 resolution were used with AGC 1 × 10^6^, MIT 200 ms, profile mode, and mass lock mass correction as above. PRM scans used a loop count of 10, 15,000 resolution, AGC target 2 × 10^5^, MIT 100 ms, NCE 26, and isolation window 2.0 m/z. The results were processed using Skyline Software and were deposited into the Panorama Public database with the unique identifier XXXX (*accession number pending*).

### 2.5. Phenotype assessment

Embryos produced by incrossing the null lines were treated in 0.2 mM PTU in system water between 6 and 20 hours post-fertilization to maintain transparency. The gross anatomy of each larva was imaged at 45 total magnification on a dissecting microscope, while one of their lenses was imaged at 200X total magnification using differential interference microscopy on an Olympus IX71 microscope. Images were taken with SPOT cameras and were cropped and oriented using ImageJ (NIH). The statistical significance in differences in proportions of defect types between samples was calculated using the “N-1” Chi-squared test (*https://www.medcalc.org/calc/comparison_of_proportions.php)* Lens diameters were measured in ImageJ with a line spanning the equator of the lens, which differs from the anterior-posterior dimension (Wang et al., 2020). Body lengths were measured from the tip of the snout to the end of the axial skeleton. Any statistically significant differences in measurements between genotypes were calculated using the *anova* and *Tukey HSD* functions in R (R. C. Team, 2020) within RStudio (Rs. Team, 2020).

Larvae were anesthetized and then transferred to 4% paraformaldehyde with 5% sucrose for storage at 4°C until cryosectioning and DAPI staining to observe the presence of lens cell nuclei. The sectioning procedure has been previously reported (Posner et al., 2013). The resulting slides with 10 micron lens sections were mounted using Vectashield Antifade Mounting Medium with DAPI (Vector Laboratories) and imaged on an Olympus IX71 microscope. DAPI intensity across the diameter of the lens was measured using the Concentric Circles plugin for ImageJ. An oval was drawn matching the lens shape, and 10 concentric circles were used to quantify the average pixel intensity along the perimeter of each circle. Any statistically significant differences in average pixel intensity between genotypes at each circle were calculated with a Mann–Whitney U test in R. All scripts and data used for this and other non-mass spectrometric analyses in this study can be found in this repository on GitHub: github.com/masonfromohio/alpha-crystallin-CRISPR-Rscripts. All lens images used in these analyses are available on Dryad.org (submission pending).

### 2.6. Quantitative PCR analysis of alpha crystallin expression

The qPCR methods were designed to meet MIQE guidelines (Bustin et al., 2009) and followed methods previously described (Posner et al., 2017). Larvae were collected at 7 dpf and stored in RNAlater (Thermo) at 4°C for up to one month until used for purification of total RNA with the Monarch Total RNA Miniprep Kit (NEB). Five larvae were pooled for each RNA purification and homogenized in DNA/RNA protection buffer with a glass homogenizer prior to proteinase K digestion, following the kit protocol. DNase treatment was performed in an RNA purification column, with a final elution in 50 µl of nuclease-free water. The concentration of collected RNA was measured in a Nanodrop One (Thermo Fisher). The ProtoScript II First Strand cDNA Synthesis Kit (NEB) was used with 600 ng of total RNA from each sample with the d(T)23 VN primer using the standard protocol in a total volume of 20 µl, and -RT controls were included.

Luna Universal qPCR Master Mix (NEB) was used to amplify 1.5 µl of each cDNA sample in a 20 µl total reaction using the manufacturer’s standard protocol on an Applied Biosystems StepOne Real-Time PCR System (Thermo). Three technical replicates were used for each cDNA sample, while two technical replicates were used with all -RT and nontemplate controls (water). Two endogenous control primer sets were included (*rpl13a* and *ef1a*). All primer sequences and previously calculated efficiencies are shown in Table 2 of Posner et al. (2017). Reactions included primers at a final concentration of 250 nm with the following parameters: hold at 95°C for 1 min; 40 cycles of 95°C for 15 s and 60°C for 30 s; fast ramp setting. Melt curve analysis was used to confirm that single products were produced, and amplification products had previously been sequenced to confirm their expected identity (Posner et al., 2017). Analysis of the resulting Cq values was identical to the process previously reported (Posner et al., 2017).

## 3. Results

### 3.1. Disruption of cryaa produced several types of lens defects that ranged in severity

We used two approaches to mutate the zebrafish *cryaa* gene to prevent the production of αA-crystallin protein. First, we injected a single guide RNA (gRNA) to produce frame shift mutations and an early stop codon (Fig. 1A). This approach successfully produced a 5 base pair deletion and early stop codon after the leucine residue at position 27 (Fig. 1B). Mass spectrometry confirmed that this mutant prevented the production of any measurable αA-crystallin protein, as shown by the absence of αA-crystallin peptide 52-65 in the *cryaa* -/- lens (Fig. 1C, right). Two additional αA-crystallin peptides that were monitored were also missing in *cryaa* -/- lenses (Supplementary Fig. S1). Differential interference contrast (DIC) microscopy of anesthetized and PTU-treated larvae at 3 days postfertilization (dpf) showed that *cryaa* homozygous null fish (cryaa^-/-^) had lens irregularities. These phenotypes included a central roughness and disorganization of central fiber cells compared to wild-type controls (Fig. 1 D-F). Because this original *cryaa* null line stopped breeding, we produced a second null line by using two gRNAs to produce an 863 bp deletion that removed a portion of the *cryaa* promoter and start codon region (Fig. 2A). This approach made it simpler to identify mutants by PCR (Fig. 2B) and reduced the potential for genetic compensation due to mutant mRNA degradation, as no truncated mRNA would be produced (El-Brolosy et al., 2019). The resulting *cryaa* null line bred well, lacked αA-crystallin peptides (Supplementary Fig. S1), and all further data presented are from these fish.

**Figure 1.**
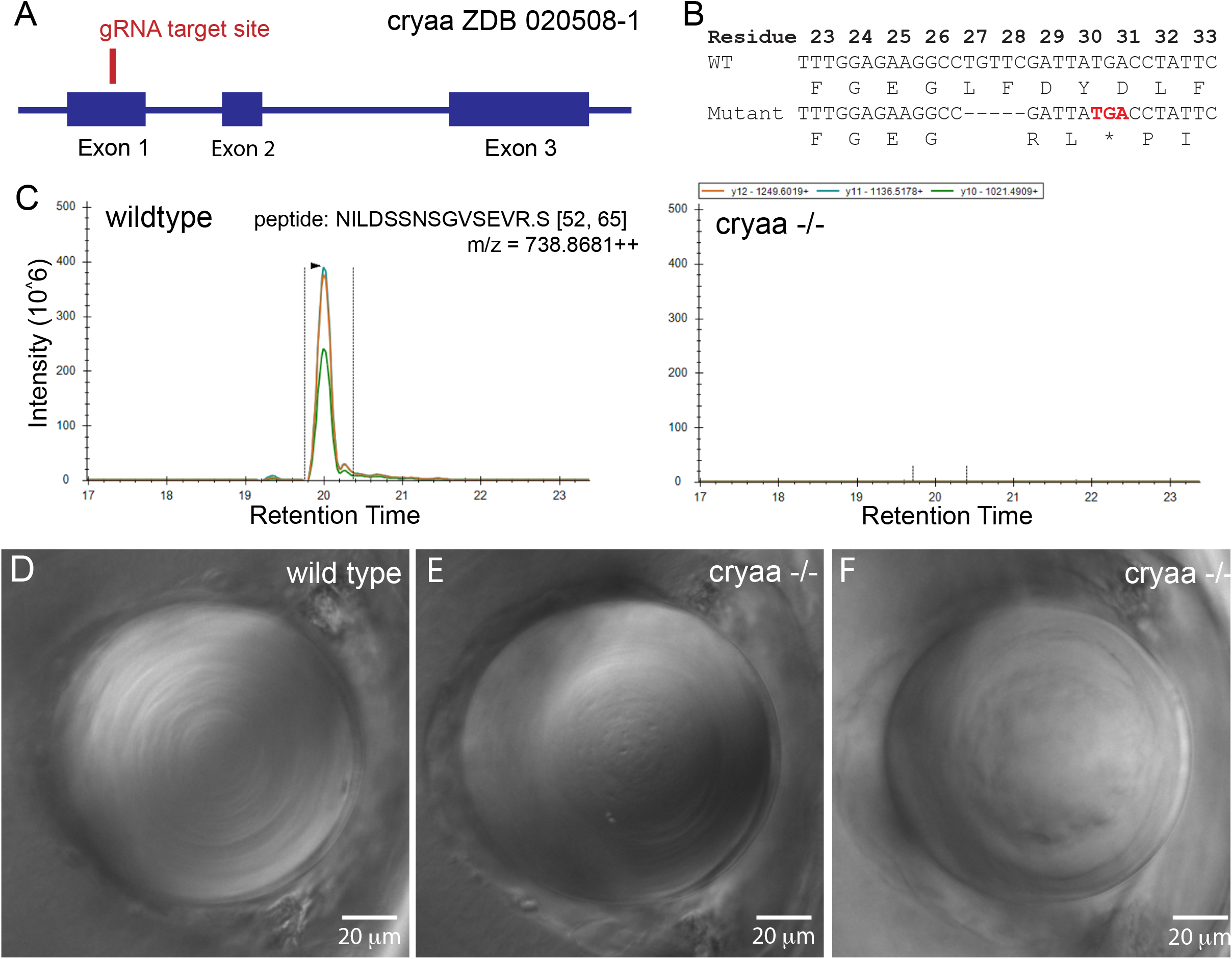
*cryaa* null mutant. Produced with a single gRNA to generate a five basepair deletion and frame shift mutation in exon 1, with the relative position noted by the red line (A). Injected embryos were outcrossed to wild-type fish to identify heterozygote mutants, with a five-base pair deletion allele identified that leads to an early stop codon after amino acid 28 (B). Incrossed heterozygotes were used to generate a homozygous null line, which was bred to assess the effects of *cryaa* loss. Mass spectrometry confirmed that embryos generated by incrossing this null line produced no detectable αA crystallin protein (C). Parallel reaction monitoring of the 3 major fragment ions of the +3 charge state of *cryaa* peptide 52-65 from the digest of wild-type lenses showed coelution of its three major fragment ions on the left and no corresponding peaks in the digest from KO lenses on the right (C). Compared to wild-type lenses at 3 dpf (C), *cryaa* null embryos showed various defects, such as roughness in the primary fiber cell region (E) and a general disorganization of central fiber cells (F). Lenses are from 3 dpf embryos, imaged using DIC optics and arranged with retina to the left. Scale bars are 20 microns.

**Figure 2.**
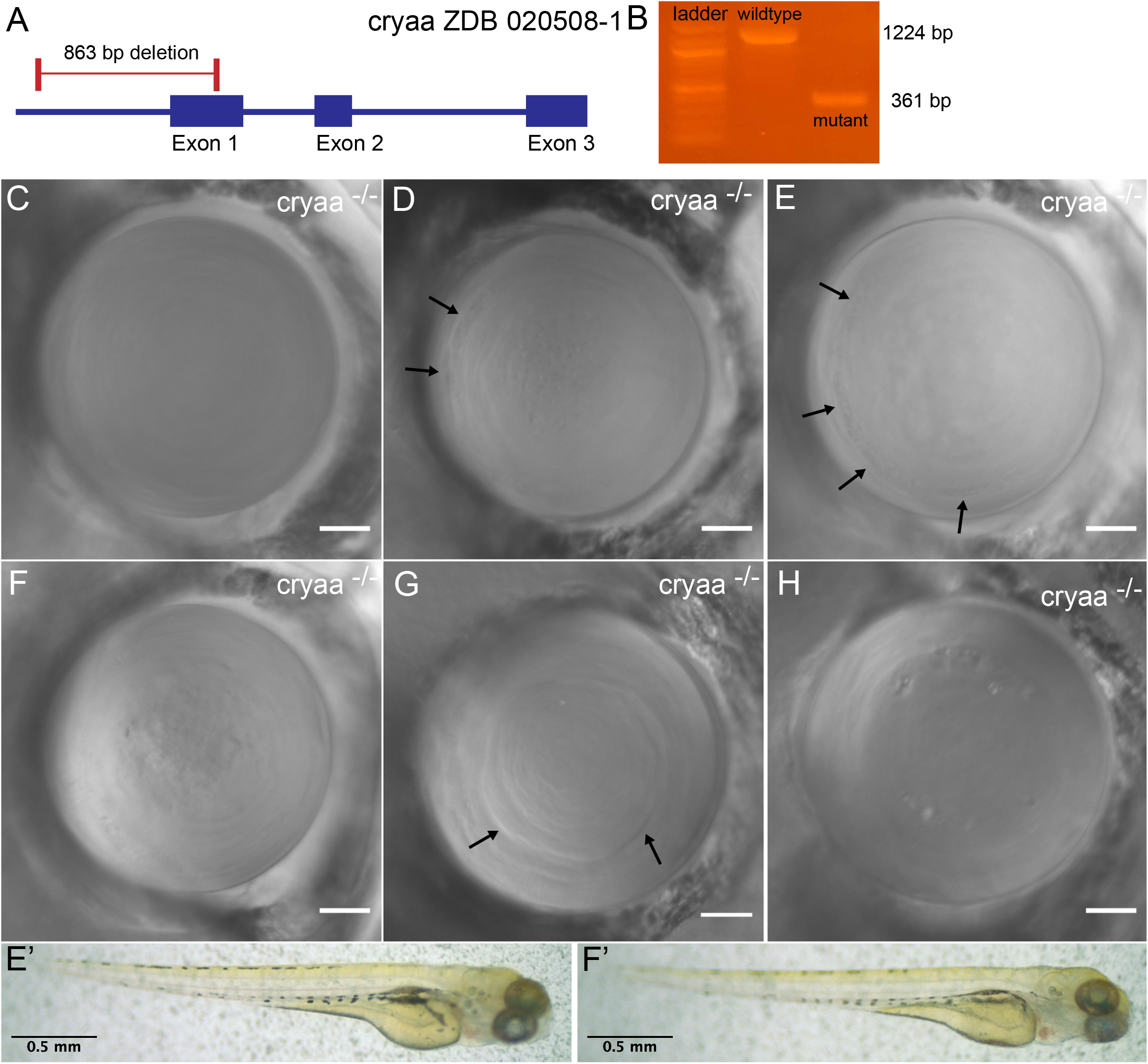
The second *cryaa* null mutant line generated by deleting the start codon and proximal promoter region produced variable lens phenotypes. Two gRNAs were designed to remove approximately 860 base pairs of the *cryaa* gene (A). This large deletion could be easily identified using PCR amplification with primers flanking the gRNA target regions (B). Some *cryaa* null embryos showed visually normal lenses by DIC imaging (C, compare to wild type lens in Fig. 1D). Typical phenotypes in abnormal lenses included roughness in central primary fiber cells (d) and general disorganization of central fiber cells (e), as seen in our single gRNA null mutant (Fig. 1). Gaps or roughness between peripheral fiber cells were also common (D and E: arrows). Some lenses combined several of these phenotypes (F). We also observed lenses with severe boundaries between fiber cells (G: arrows) and pitting (H). Lenses shown are either 3 or 4 dpf and arranged with retina to the left. Scale bars are 20 microns. Matching embryos for lenses E and F are shown to indicate a lack of general morphological defects (E’ and F’).

A substantial proportion of *cryaa* null larvae had lenses that appeared similar to those of wild-type fish (Fig. 2C). Others showed phenotypes that we classified into five categories. Some lenses had central roughness that appeared as many small darker spots in the region of the primary fiber cells (Fig. 2D). Some lenses contained elongated roughened areas that appeared to trace the edges of peripheral fiber cell boundaries (Fig. 2 D and E, black arrows). We also found lenses with a disorganization of central fiber cells (Fig. 2 E and F). These lenses did not show regular circular rings of central fiber cells. Some lenses showed more severe boundaries between fiber cells, possibly at the interface between the lens nucleus and cortex (Fig. 2G, black arrows), and some lenses exhibited large pitting (Fig. 2H).

### 3.2. Frame shift mutations of the two paralogs for zebrafish αB-crystallin genes did not produce significant overall lens defects

Single gRNAs were designed to target exon 1 in both the *cryaba* and *cryabb* genes (Fig. 3). Injected embryos were raised, and those with mutations leading to frameshifts and early stop codons were identified. A single basepair deletion in *cryaba* led to an altered amino acid sequence after residue 14 and a stop codon at residue 65 (Fig. 3B). Mass spectrometry indicated that lenses from *cryaba* mutant adults expressed no detectable αBa-crystallin, as evidenced by missing tryptic peptide 80-89 in the mutant lens (Fig. 3C, right). An 8 bp deletion in *cryabb* produced altered amino acids after residue 9 and a stop codon at residue 26 (Fig. 3D & E). Digests of adult *cryabb* homozygous mutant fish lenses had undetectable levels of peptide 12-20 (Fig. 3F, right). However, of the three peptides monitored in *cryabb* adult mutant lenses, the most C-terminal peptide (residues 82-92) was still detected (Supplementary Fig. S1). The *cryabb* mutant has a start codon in frame that could direct production of a partial peptide including residues 59 through 165, which is consistent with our mass spectrometry data.

**Figure 3.**
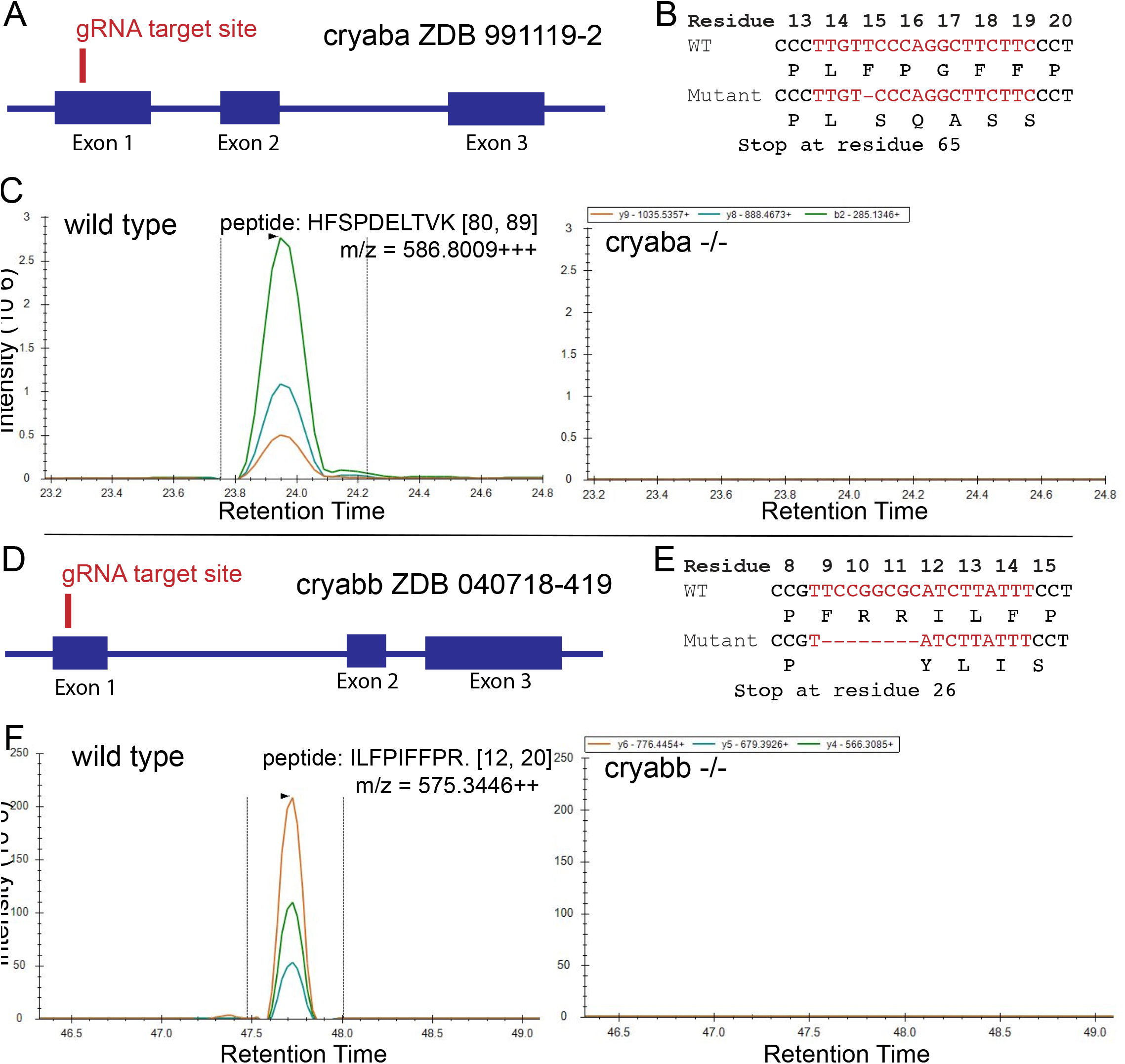
Production of *cryaba* and *cryabb* frameshift mutants. Single gRNAs were designed to target the first exons of the zebrafish *cryaba* (A) and *cryabb* (D) genes. After injection into wild-type zygotes, founders were outcrossed to wild-type fish, and resulting heterozygotes with deletions that caused frameshift mutations leading to early stop codons were detected (B and E). Mass spectrometry analysis of adult lenses from mutant lines confirmed the loss of protein coded by each gene. Parallel reaction monitoring of the +3 charge state of αBa-crystallin tryptic peptide 80-89 from the digest of wild-type lenses detected coelution of its three major fragment ions (left), and these were not detectable in *cryaba* mutant adult lenses (C). Similarly, the +2 charge state of αBb-crystallin tryptic peptide 12-20 from the digest of wild-type lenses detected coelution of its three major fragment ions (left), and these were missing in digests from *cryabb* mutant adult lenses on the right (F).

The prevalence of lens defects in larvae generated by our *cryaa, cryaba* and *cryabb* mutant lines was quantified at 3 and 4 dpf to assess the impacts of loss of each α-crystallin on lens development (Fig. 4). Lens defects were statistically significantly increased in mutant *cryaa* larvae at both 3 and 4 dpf and in *cryaa* crispant larvae at 4 dpf compared to wild-type larvae. There was no significant increase in overall lens defects in the *cryaba* or *cryabb* mutant larvae. The appearance of central roughness was characteristic of our *cryaa* null larvae and not significant in any other sample. Peripheral fiber cell defects and severe fiber cell boundaries were significantly increased at some of our *cryaa* null and crispant timepoints. While lower in proportion, these fiber cell defects were significantly higher at some *cryaba* and *cryabb* mutant ages compared to wild-type larvae. Central disorganization of the lens was only significantly increased in 4 dpf *cryaa* null larvae. Pitting was the most common defect seen in wild-type larvae and not significantly increased in any of our mutant larvae.

**Figure 4.**
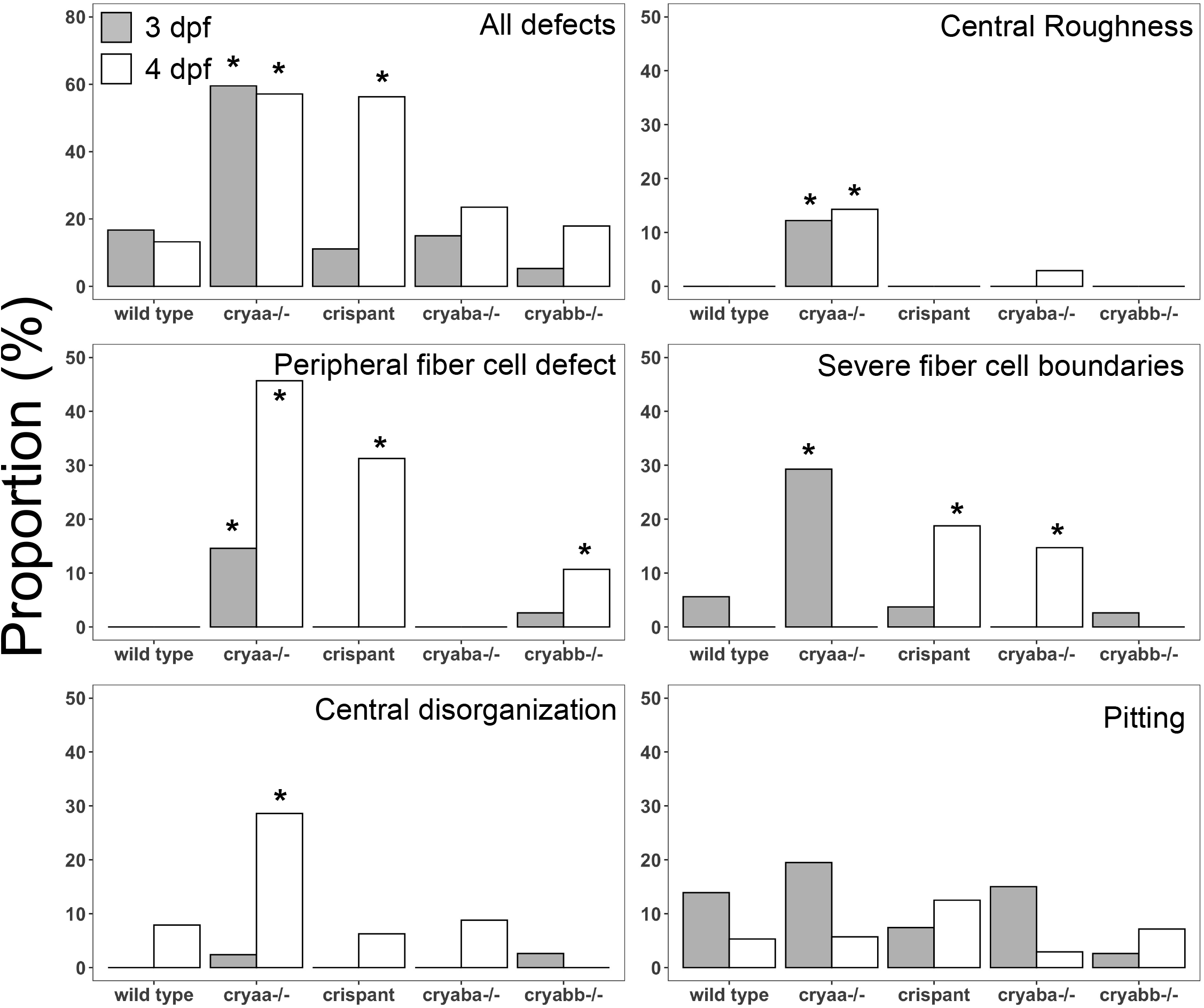
Prevalence of different lens abnormalities in wild-type, α-crystallin mutant and *cryaa* gRNA-injected larvae at 3 and 4 dpf. Each graph shows the proportion of larvae with that defect type. Larvae examined at 3 and 4 dpf were wild-type (n= 36/38), offspring of homozygous mutants for *cryaa* (n=41/35), *cryaba* (n=40/34) and *cryabb* (n=38/28), and embryos injected with a four-gRNA mix targeting the *cryaa* gene (crispants; n=27/16). Asterisks indicate statistically significant differences from age-matched wild-type larvae (“N-1” Chi-squared test).

All larvae used to assess lens phenotype were measured to record body length and lens diameter at 3 and 4 dpf (Fig. 5). These data show an increase in body length from 3 to 4 dpf in all genotypes, as would be expected. Body length was more varied between two independent wild-type samples than between wild-type and mutant samples, suggesting that gene knockout did not alter the overall fish growth rate compared to wild-type fish. There appeared to be little to no difference in lens diameter between the 3 and 4 dpf samples, which produced a decrease in the lens to body length ratio between those two timepoints for all genotypes. The only measurement that was significantly different from both wild-type samples was a reduced lens diameter of *cryabb* mutant larvae at 4 dpf.

**Figure 5.**
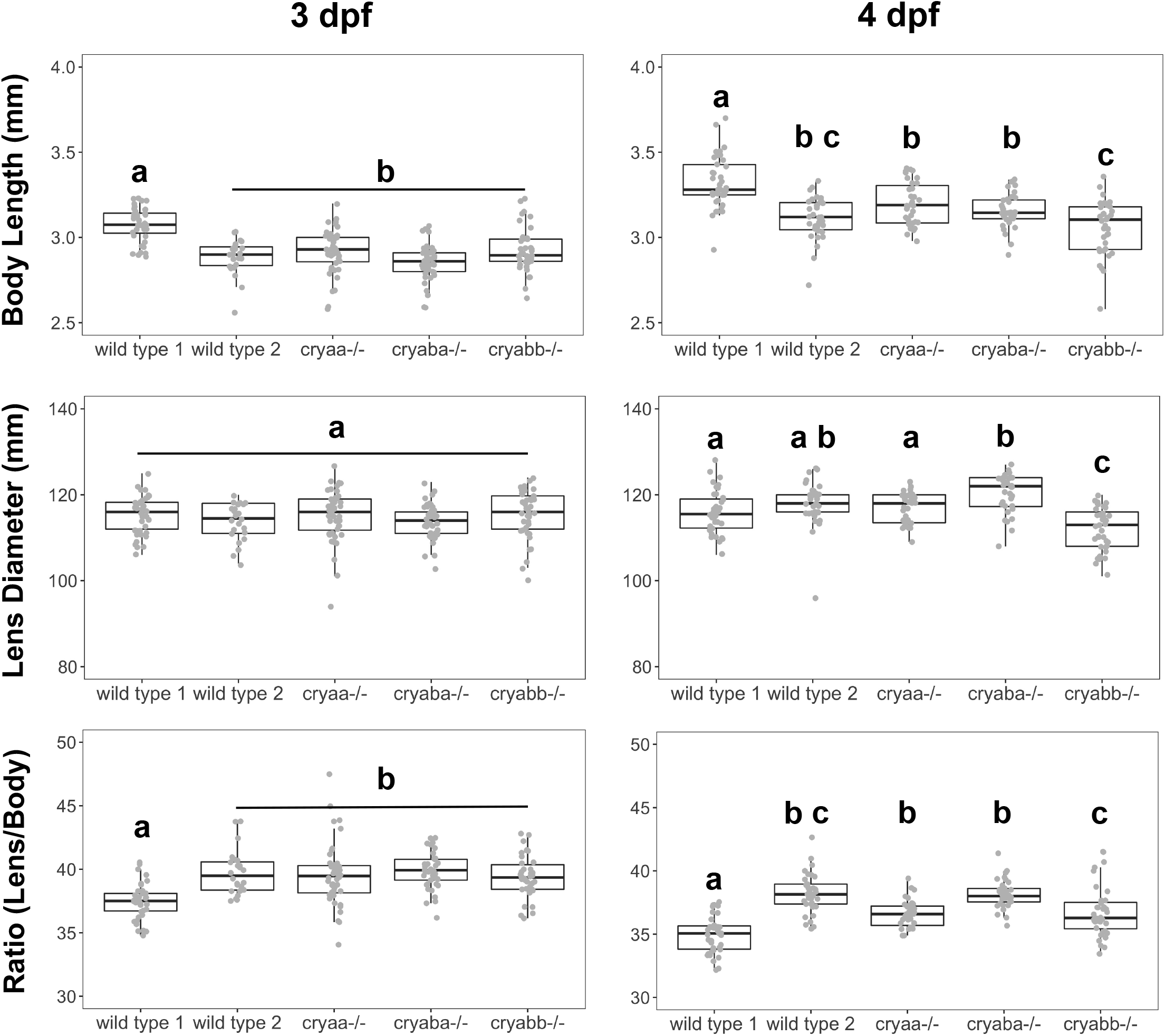
Comparison of lens diameters and body lengths between wild-type and α-crystallin knockout embryos. Box and whisker plots show measurements at 3 and 4 days post-fertilization (dpf). Analysis of variance with Tukey’s post-test was used to identify statistically significant differences between groups. Letters indicate statistical groups within each panel.

### 3.3. Loss of α-crystallins did not disrupt fiber cell denucleation

Zebrafish lens fiber cells elongate and lose their nuclei between 2 and 3 dpf. We examined lenses histologically at 3 and 4 dpf to determine if this process was altered by loss of each α-crystallin (Fig. 6). Wild-type lenses showed that while the timing of this process can vary, the majority of lenses cleared all nuclei from central fiber cells by 3 dpf, leaving elongated nuclei at the posterior margin (Fig. 6A, white boxes). We consistently found two posterior zones of extended fiber cells bordering a denucleated space between them (Fig. 6A, asterisk), with what appeared to be a single layer of nuclei adjacent to the retinal neurons. Lenses contained rings of DAPI staining just anterior to the remaining, elongated fiber cells and extending forward with a gap between the anterior margin and the epithelial fiber cells (Fig. 6A, white arrows). The posterior-most gap between elongated fiber cells was a good indicator that sections imaged were through the equator of the lens. Elongated fiber cell nuclei in wild-type lenses ranged from flat to wavy and occasionally retained an oval shape (Fig. 6C, white arrow). At 4 dpf, the posterior fiber cell nuclei stacks appeared thinner, and DAPI rings were less common.

**Figure 6.**
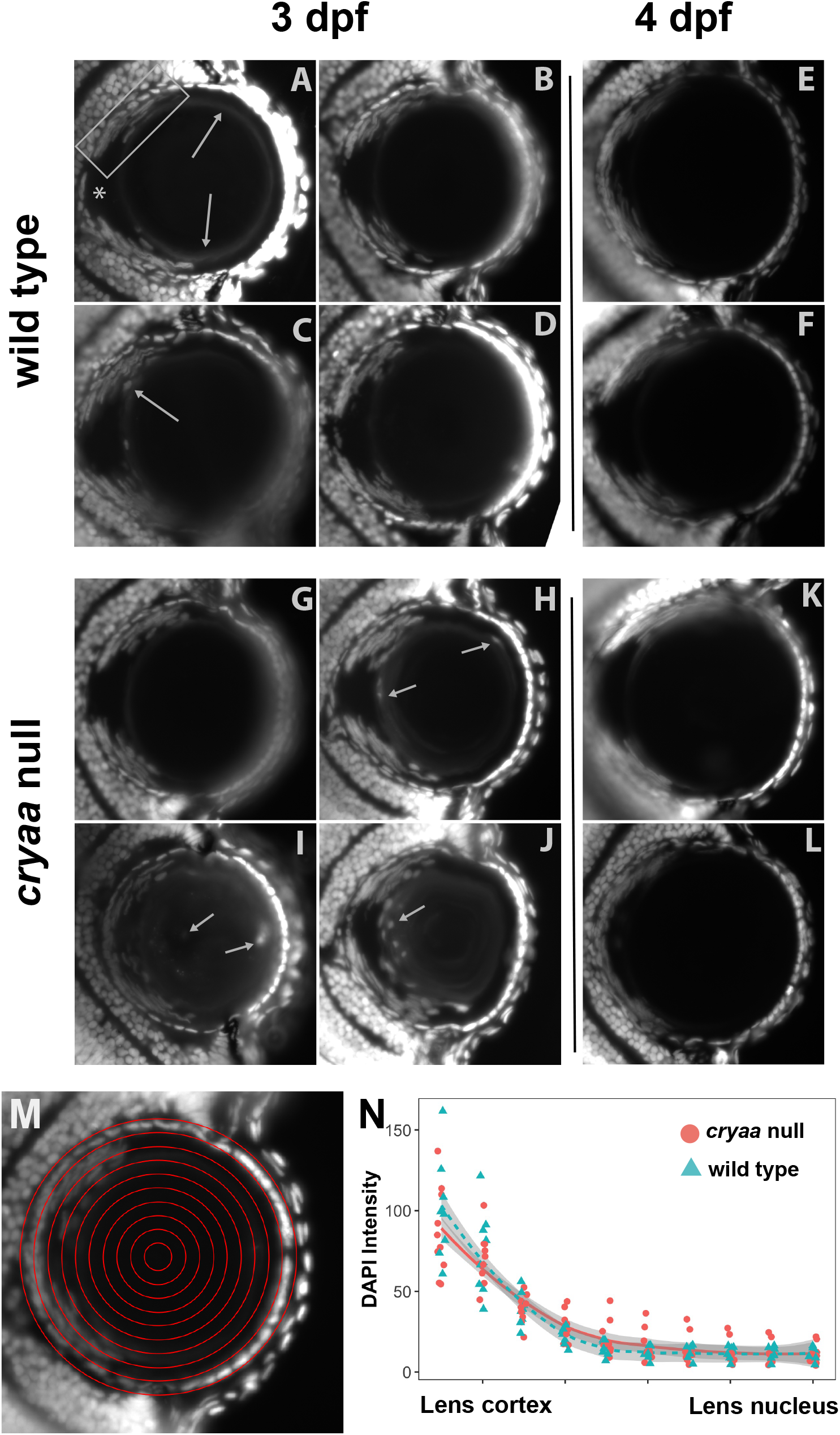
Cryosectioned lenses at 3 and 4 dpf were stained with DAPI to assess the clearance of fiber cell nuclei. Four representative lenses are shown for the wild-type (A-D) and *cryaa* null lines (G-J) at 3 dpf, and two representative lenses are shown at 4 dpf (E-F; K-L). Most images were taken at 200X total magnification, with some taken at 100X to capture more lens cell nuclei in one focal plane. The retina is oriented to the left in each image, with the cornea to the right. The four images at 3 dpf for each genotype demonstrate the range of phenotypes found, with the top left panel the most normal and bottom right the most abnormal. The rectangle in panel A highlights one of two regions of flattened, extended fiber cell nuclei at the retinal side of the lens, separated by a nucleus-free zone at *. Arrows indicate details described in the results section. Concentric circles were used to measure the average pixel density of DAPI staining across the lens (see M as an example). While some *cryaa* null lenses showed higher DAPI intensity in inner areas of the lens at 3 dpf (N), these differences were not statistically significant (Mann–Whitney U test).

The lenses of *cryaa* null 3 dpf larvae also varied in their DAPI staining. However, more abnormal phenotypes were seen. A small number of *cryaa* null lenses retained fiber cell nuclei deep to the central DAPI staining ring (Fig. 6 I and J) or on the ring itself (Fig. 6H). Quantitation of pixel intensity also showed that some *cryaa* null larval lenses had higher amounts of DAPI staining material in the deeper parts of the cortex and lens nucleus, but the mean pixel intensity did not differ statistically from that of wild-type larvae (Fig. 6N). None of the 4 dpf *cryaa* null larvae examined retained abnormal fiber cell nuclei. None of the *cryaba* or *cryabb* mutant 3 dpf larvae examined showed retained fiber cell nuclei within the central lens (Supplementary Fig. S2). However, some lenses did appear to have thicker fiber cell nuclei stacks and possibly more nuclei with oval shape, although these features were also seen in some wild-type lenses as described above.

### 3.4. CRISPR deletion of each α-crystallin gene did not alter the expression of the other two

Quantitative PCR (qPCR) was used to measure α-crystallin gene expression levels in wild-type and our mutant lines. This allowed us to determine whether mutation led to a decrease in mRNA levels for the damaged gene or changes in expression of the other α-crystallin genes. Whole larvae were assessed at 7 dpf, as earlier work showed low levels of *cryaba* and *cryabb* expression through 4 and 5 dpf, respectively (Posner et al., 2017), and almost no expression in the lens (Farnsworth et al., 2021). We found reduced levels of *cryaa* mRNA in 7 dpf homozygous *cryaa* mutants produced by deleting the proximal promoter and start codon (Fig. 7A). The expression levels of *cryaa* did not change in *cryaba* or *cryabb* mutant larvae compared to wild-type larvae (Fig. 7A). The *cryaba* and *cryabb* mutant lines were not produced by deleting the promoter and start codon but instead were produced with a single gRNA producing frame shift mutations and early stop codons (Fig. 3). *cryaba* mRNA levels were not different in the *cryaba* mutant larvae compared to wild-type larvae, nor were they increased in the other mutant lines (Fig. 7B). Messenger RNA for *cryabb* was reduced in *cryabb* mutant larvae compared to other mutant lines (Fig. 7C). While the delta Cq values for *cryabb* mRNA in *cryabb* mutants were all lower than those in wild-type larvae, this difference was not statistically significant. Messenger RNA levels for *cryabb* were not significantly increased in *cryaa* or *cryaba* mutants compared to wild-type larvae.

**Figure 7.**
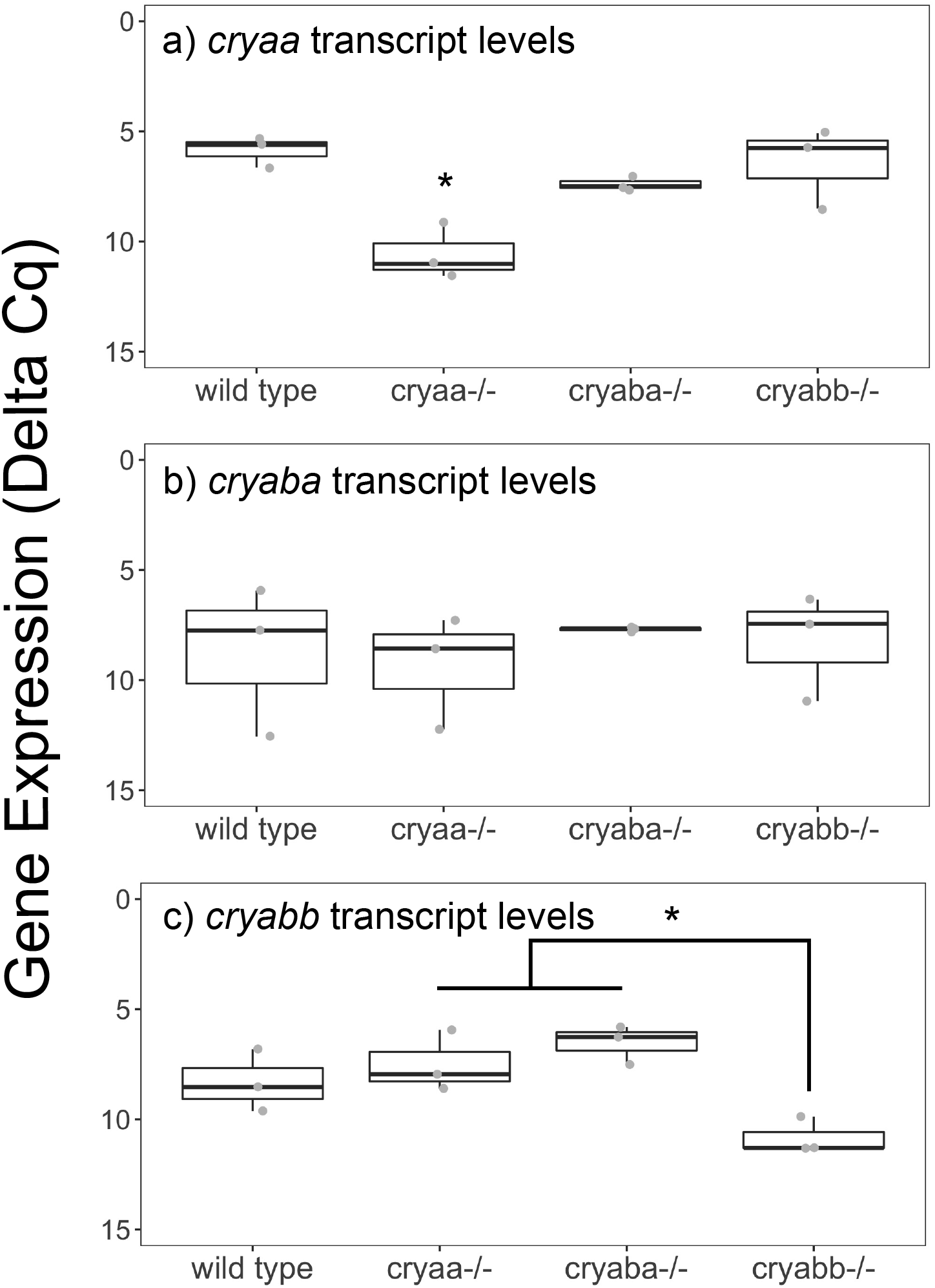
Expression of each α-crystallin gene assayed by qPCR in whole bodies of wild-type and mutant larvae at 7 dpf. Box and whisker plots show delta Cq values generated with primers for *cryaa* (A), *cryaba* (B) and *cryabb* (C) in three biological replicates of the genotypes indicated on the x-axes. Y-axes are inverted, as lower delta Cq values indicate greater expression. Asterisks indicate statistically significant differences based on ANOVA with Tukey’s post-hoc test (*p value* < 0.05).

## 4. Discussion

We produced mutant zebrafish lines for each of the three α-crystallin proteins. Lack of αA-crystallin caused a variety of visible lens defects in mutant larvae not seen or present at lower frequency in wild-type larvae. Mutation of either αB-crystallin did not cause an increase in overall lens abnormalities, although our *cryabb*-null line expressed a truncated αBb-crystallin. Loss of αA-crystallin did not alter overall lens growth or significantly disrupt the fiber cell denucleation important for lens development. Finally, none of the gene mutations produced compensatory changes in the mRNA levels of the others. These data suggest that loss of αA-crystallin makes zebrafish lenses more susceptible to defects in development but does not always trigger these defects. Loss of either αBa-crystallin or a full-length αBb-crystallin produced little significant change in the lens through four days of development.

The observation of lens abnormalities in our CRISPR-produced *cryaa* null larvae agrees with previous work done with a TALEN-produced *cryaa* knockout line (Zou et al., 2015). However, while lens images from that study showed central roughness and pitting of the lens, the authors did not indicate the central disorganization, peripheral fiber cell roughness or irregular fiber cell boundaries identified in our *cryaa* null larvae. The proportion of null incross larvae with lens defects in the TALEN study was approximately 85%, slightly more than our finding of 60%. That study separated their phenotypes into “minor” and “major”, with just over 40% of larvae having minor lens defects. Despite the different methods used to characterize lens defects in our present study and this past TALEN work, it seems that the observed penetrance of *cryaa* loss is comparable. It is notable that the effects of αA-crystallin loss can range from no apparent defect in lens structure to a variety of defects that can occur singly or in combination. A similar range of lens effects was seen after CRISPR knockout of the zebrafish aquaporin 0 paralogs, *mipa* and *mipb* (Vorontsova et al., 2018). Our results suggest that αA-crystallin is not essential for proper lens development through 4 days post-fertilization but that its loss predisposes the lens to defects. The only other model species with an αA-crystallin knockout line is the mouse (Brady et al., 1997). Due to the *in utero* development of the mouse and its pigmentation, it was likely not feasible to make similar observations of lens development at comparable stages to this study.

However, that study showed that *cryaa*-null mice develop cataracts by 7 weeks of age, indicating the importance of the protein in lens biology. The impact of *cryaa* loss on zebrafish lens clarity as animals age can be addressed in the future using our validated lines. The impact of CRISPR-generated null mutation in *cryaa*, reported here, can be compared to prior knockdowns by morpholino injection. We previously reported a lack of lens defects in zebrafish larval lenses at 3 dpf after using a morpholino to block the translation of αA-crystallin (Posner et al., 2013). Similar results were reported in a later study (Hinaux et al., 2014). Neither study imaged lenses using DIC microscopy, so it is possible that defects such as those identified in the current study were missed. It is also possible that small amounts of αA-crystallin produced during those morpholino experiments were sufficient to prevent the defects found in this study.

Our data do not directly address the mechanisms behind the lens defects seen in our *cryaa* null lenses. Histology and DAPI staining of 3 and 4 dpf larvae indicate that while there may be some delay in fiber cell denucleation in *cryaa* null lenses, this delay was only found in a small proportion of lenses at 3 dpf and gone by 4 dpf. Because αA-crystallin has documented functions as an anti-aggregation chaperone that can buffer the effects of stress in the lens (Horwitz, 1992), it is possible that its loss may make the lens more susceptible to protein aggregation. This likely led to the cataract seen in *cryaa* null mouse lenses (Brady et al., 1997), and since zebrafish αA-crystallin has chaperone activity (Dahlman et al., 2005; Koteiche et al., 2015), aggregates in central fiber cells may lead to observed central roughness. This hypothesis could be tested by future proteomic analysis of insoluble proteins. Other defects we observed hint at a breakdown in normal fiber cell arrangement. Under DIC imaging, the fiber cell borders in the zebrafish lens were visible as regular concentric rings (Fig. 1D). Our observation of central disorganization suggests defects in the arrangement of central fiber cells. A separate defect appeared as short regions of rough, expanded space between peripheral fiber cells. These three anatomical defects were visually distinct and may stem from different mechanistic defects in fiber cell interactions. It has been proposed that αA-crystallin plays a direct role in fiber cell differentiation (Boyle and Takemoto, 2000). While our *cryaa* null larvae did not show extended defects in fiber cell nucleus degradation, there may be other aspects of fiber cell differentiation impacted by αA-crystallin loss.

Our finding that lens diameters in *cryaa* null lenses at 3 and 4 dpf were similar to wild-type larvae matches the results of a previous knockout study (Zou et al., 2015). Our *cryaba* and *cryabb* mutant larvae also appeared similar in body length and lens diameter to wild-type larvae. We included two separate samples of wild-type larvae to assess possible variation between breedings. The statistically significant variation in body length between these two wild-type samples suggests that variation between groups can be high and should be taken into account when looking for changes in lens growth resulting from gene knockout. The observation that lens diameter was essentially unchanged between 3 and 4 dpf, while body length increased, was unexpected considering that past studies have found linear relationships between lens diameter and body length, although these studies did not examine fish prior to 7 dpf (Collery et al., 2014; Wang et al., 2020). It is possible that in these earlier stages, body length and lens diameter growth are not parallel.

We found little evidence that mutation of either αB-crystallin gene in zebrafish caused abnormalities in lens development or structure. Neither our *cryaba* nor *cryabb* mutant larvae showed a statistically significant increase in the proportion of all lens defects compared to wild-type larvae (Fig. 4). Fiber cell denucleation also appeared normal in both αB-crystallin mutant lines. The lack of developmental lens defects in these fish is consistent with knockout studies in mice in which loss of the single αB-crystallin gene had no effect on lens clarity through 39 weeks of age, although early developmental stages were not examined (Brady et al., 2001). However, previously published CRISPR mutants for *cryaba* and *cryabb* did produce lens defects at 4 dpf in a large proportion of larvae (Mishra et al., 2018). In that study, approximately 75% and 50% of *cryaba-* and *cryabb*-null larvae, respectively, showed lens abnormalities. Defects were categorized as “minor” and “major” with little description of defect type. Two published images of lenses from a *cryaba* heterozygote cross show roughness or pitting as well as some gaps between fiber cells. Previous work by the same lab identified far more severe lens defects in zebrafish larvae after injection of translation-blocking morpholinos targeting *cryaba* and *cryabb* (Zou et al., 2015). However, general gross defects in the resulting larvae suggest general toxicity from morpholinos and not a lens-specific effect. In their more recent CRISPR lines, there were no reported general larval defects, further suggesting that the morpholino-induced lens defects were not specifically due to the loss of αB-crystallin proteins. While we found some significant increase in fiber cell defects in our *cryaba* and *cryabb* mutant larvae compared to wild-type larvae, these proportions were far lower than those reported in past studies.

Existing data suggest that neither *cryaba* nor *cryabb* are substantially expressed in the zebrafish lens through 5 dpf based on shotgun proteomics (Greiling et al., 2009) and single-cell RNA-Seq (Farnsworth et al., 2021). These published findings fit with the lack of significant early lens phenotypes in our αB-crystallin mutant larvae. The reason for the discrepancy in our present CRISPR work and a past study identifying lens defects in *cryaba* and *cryabb* knockout zebrafish (Mishra et al., 2018) is unclear. Our mass spectrometry data indicated a total loss of all detectable αBa-crystallin peptides in *cryaba* null adult lenses, indicating that our *cryaba* null line is incapable of producing αBa-crystallin protein.

However, the presence of the αBb-crystallin peptide 82-92 in adult lenses from our *cryabb* null line suggest that these fish are capable of producing a truncated protein likely spanning residues 59-165. This truncated C-terminal αBb-crystallin would contain the anti-aggregation chaperone site identified in mammalian αB-crystallin (Bhattacharyya et al., 2006). Since this αBb-crystallin peptide was identified by analyzing adult lenses our data do not indicate that this partial αBb-crystallin protein is produced in larval lenses at the ages analyzed in this study (3 and 4 dpf). The production of a partial protein even after a frame-shift mutation highlights the value in using mass spectrometry to confirm the total loss of expression from a CRISPR targeted gene.

The variety of approaches used to produce αA-crystallin mutants in this study allows us to compare their utility for studying resulting phenotypes. The quickest way to screen genes for their role in development is to directly examine larvae injected at the zygote state (crispants). There is good evidence that this approach is effective when injecting a mix of four gRNAs (Wu et al., 2018). In our own hands, we found this approach effective when targeting genes already known to play an essential role in lens development. For example, injection of a four-gRNA mix targeting the transcription factor *foxe3* closely replicated phenotypes reported in a stable zebrafish null line (Supplementary Fig. S3; (Krall and Lachke, 2018)). However, when targeting *cryaa*, this approach did not produce similar levels of phenotypes seen in our null line (Fig. 4). The proportion of each defect differed between the two approaches, and the levels of some defects also differed between 3 and 4 dpf. These variations in phenotype could be due to the mosaic nature of gRNA-injected larvae, in which some lens cells contain mutant *cryaa* alleles and others contain wild-type alleles. The expression of αA-crystallin in unaffected cells may be enough to prevent the defects seen in our null line. We previously showed that adult zebrafish lenses produce less αA-crystallin than mammalian lenses (Posner et al., 2008), and our past qPCR and single-cell RNA-Seq data suggest that lens *cryaa* expression through 5 dpf is relatively low as well (Farnsworth et al., 2021; Posner et al., 2017). Since the complete loss of αA-crystallin in our *cryaa* null line only led to visible defects in 60% of larvae, the expression of reduced amounts in our injected embryos may be sufficient to suppress the phenotype. It is also possible that differences are simply due to the stochastic nature of phenotype development. Considering the dynamic changes occurring in the zebrafish lens between 3 and 4 dpf, it is best to quantify phenotypes for each day separately in future studies.

Because our initial *cryaa* null line generated with a single gRNA and containing a frameshift mutation stopped breeding, we took the opportunity to generate a second null line with a large deletion removing the proximal promoter region and start codon. Both lines generated similar lens defects, reinforcing the conclusion that these defects resulted from the loss of αA-crystallin and not off-target effects. There are two other benefits to generating mutant knockout alleles by deletion. First, these deletion alleles are simpler to screen, as they are identifiable by PCR without subsequent sequencing. Second, deletion of the promoter and start codon reduces the chance of possible genetic compensation after nonsense-mediated mRNA decay (El-Brolosy et al., 2019). The loss of the targeted gene’s mRNA can also be monitored by qPCR (Fig. 7). Our two αB-crystallin mutant lines were produced with single gRNAs and frameshift mutations. While our proteomics work analyzed the loss of each protein in our null adult fish lenses, qPCR showed a reduction in *cryabb* mRNA in the *cryabb* null line but no reduction in *cryaba* mRNA in the *cryaba* null line. This difference may reflect the unpredictability of nonsense-mediated decay and shows that mRNA levels do not necessarily reflect eventual protein levels. It is also possible that this difference in mRNA levels reflects primer location, as our *cryaba* qPCR primers annealed within exon 1, just downstream of the gRNA target site, while our *cryabb* primers annealed to the 5’ UTR.

Several key questions remain unanswered in this study. It is unclear how the loss of αA-crystallin leads to the visual defects observed by DIC microscopy. Lenses from mutant α-crystallin mice have been analyzed by proteomics and RNA-Seq to identify changes in global gene expression (Andley et al., 2018, 2013). A similar approach would be valuable with our zebrafish null lines. We have only presented data through 7 dpf. It will be interesting to see how αA-crystallin loss impacts lens clarity and optics as fish age. While we know that both αB-crystallin genes are expressed in the adult lens, we do not know at what age this expression begins and how αB-crystallin loss may impact lenses as they age. This question is particularly interesting, as *cryaba* is lens specific in adults, while *cryabb* is expressed broadly, suggesting an ontogenetic shift in the function of these genes and their proteins. Why *cryaba* is expressed outside the lens in early development and whether it or *cryabb* plays an important developmental role in those extralenticular tissues are also unknown. Answers to these questions would add to the understanding of why these small heat shock protein genes were coopted as key players during vertebrate lens evolution.

## Acknowledgments

We would like to thank the many members of the zebrafish research community who were helpful with technical suggestions. The plasmid-free protocol used to generate gRNAs was shared with us by Dr. Cody Smith at the University of Notre Dame. Dr. Jennifer Phillips at the University of Oregon first suggested our double gRNA deletion approach for generating our *cryaa* mutant. Caitlin Puff contributed to the measurements of larval body length and lens diameters. This work was supported by the NIH National Eye Institute (R15 EY13535) to MP, by the Provost Office at Ashland University (AU) and by the AU Choose Ohio First program that provided summer research support to KM. Mass spectrometric analysis was supported by the NIH National Institute (P30 EY010572).

## Author Contributions

MP conceived and designed the study. KZ and LLD designed and conducted the proteomic experiments. MP, KLM, BA, SB, AR, KF, AH, TK, EK, MSM and TS contributed to the production of CRISPR null lines and/or analyzed the resulting phenotypes. MP, KLM, AR, AH, TK, and LLD wrote the manuscript. All authors reviewed the manuscript.

## Data Availability

Data spreadsheets and R scripts used for Figures 4-7 can be found in the following GitHub repository: github.com/masonfromohio/alpha-crystallin-CRISPR-Rscripts. DIC images and DAPI images of lenses will be made available on Dryad.org. Skyline analysis of mass spectrometry data will be made available on PanoramaWeb (panoramaweb.org)

