## Supplemental Figure 1 for "Effects of α-crystallin gene knockout on zebrafish lens development"

**Legend Supplementary Fig. S1.**

Chromatographic traces of three tryptic peptides from each of αA, αBa, and αBb-crystallins from wild-type and *cryaa*, *cryaba*, and *cryabb* mutant lenses from adults. Wild-type and *cryaba* mutant lenses were 12 months old and *cryaa* and *cryabb* mutant lenses were 6 months old. **A-C**: αA 53-66, 90-100, and 121-146, respectively. **D-F**: αBa 80-80, 90-100, and 155-168, respectively. **G-I:** αBb: 12-20, 46-58, and 82-92, respectively. In each panel A-I: 1: wild-type lens 1, 2: wild-type lens 2, 3: *cryaa* mutant lens (first generated mutant), 4: *cryaa* null lens (second generated mutant), 5: *cryaba* mutant lens 1, 6: *cryaba* mutnat lens 2, 7: *cryabb* mutant lens 1, 8: *cryabb* mutant lens 2. In each pair of chromatograms, the upper trace shows the peak for the designated peptide precursor ion of the designated charge state, and the lower trace shows three corresponding coeluting peptide y- and b- fragment ions. The intensity scales for each grid of chromatographic peaks were not set at the same scale so that lower intensity peaks could be observed. When peaks were not observable in digests of null lenses, the intensity scale was set at the same value as the lowest intensity scale on chromatograms from lenses where the respective α-crystallin was not knocked out. Underlined cysteines denote alkylation by iodoacetamide. Chromatograms were generated using Skyline software and can be accessed on PanoramaWeb (accession number pending).

A) αA: R.NILDSSNSGVSEVR.S [53, 66]

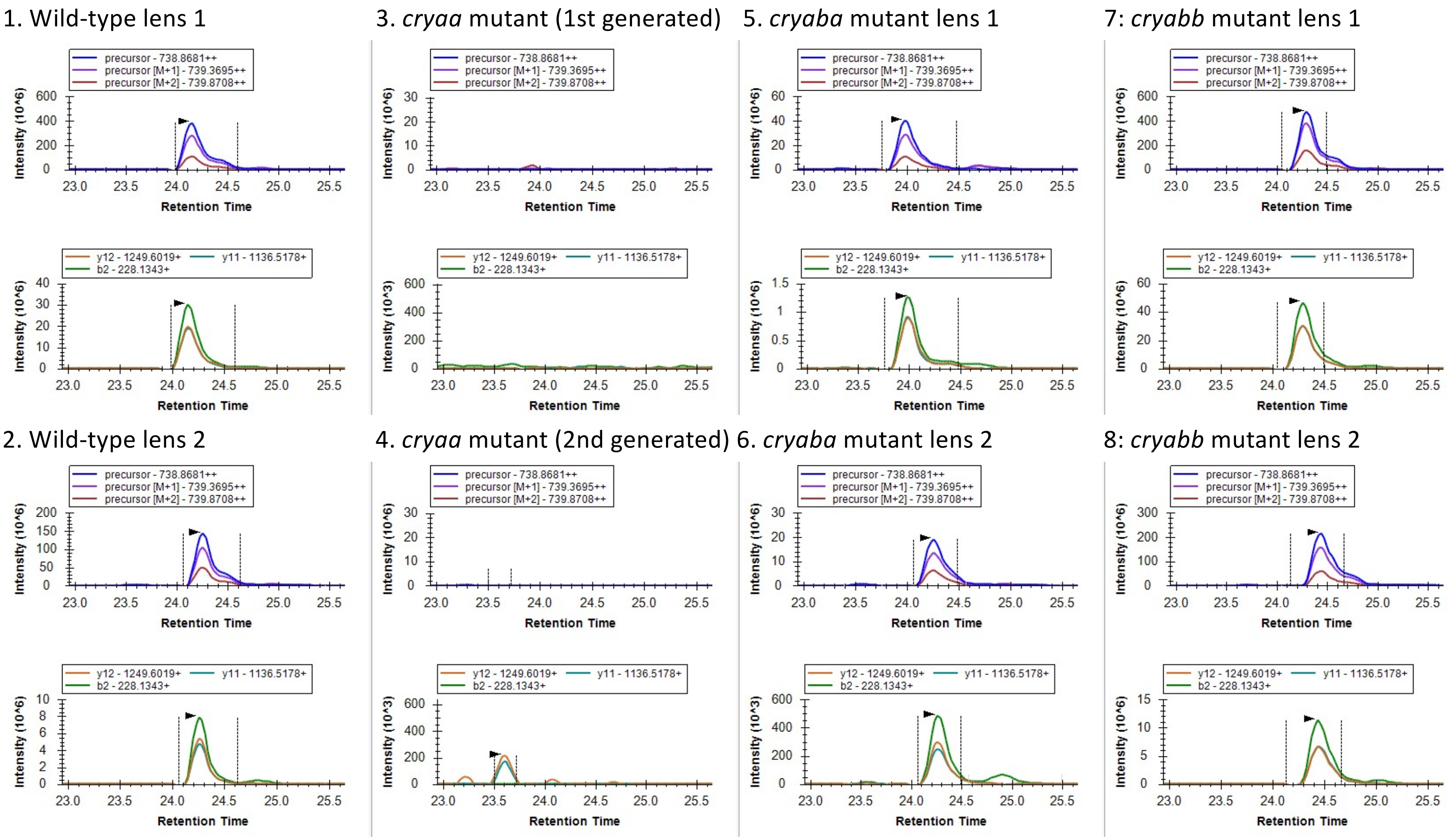

B) αA:K.VTDDYVEIQGK.H [90, 100]

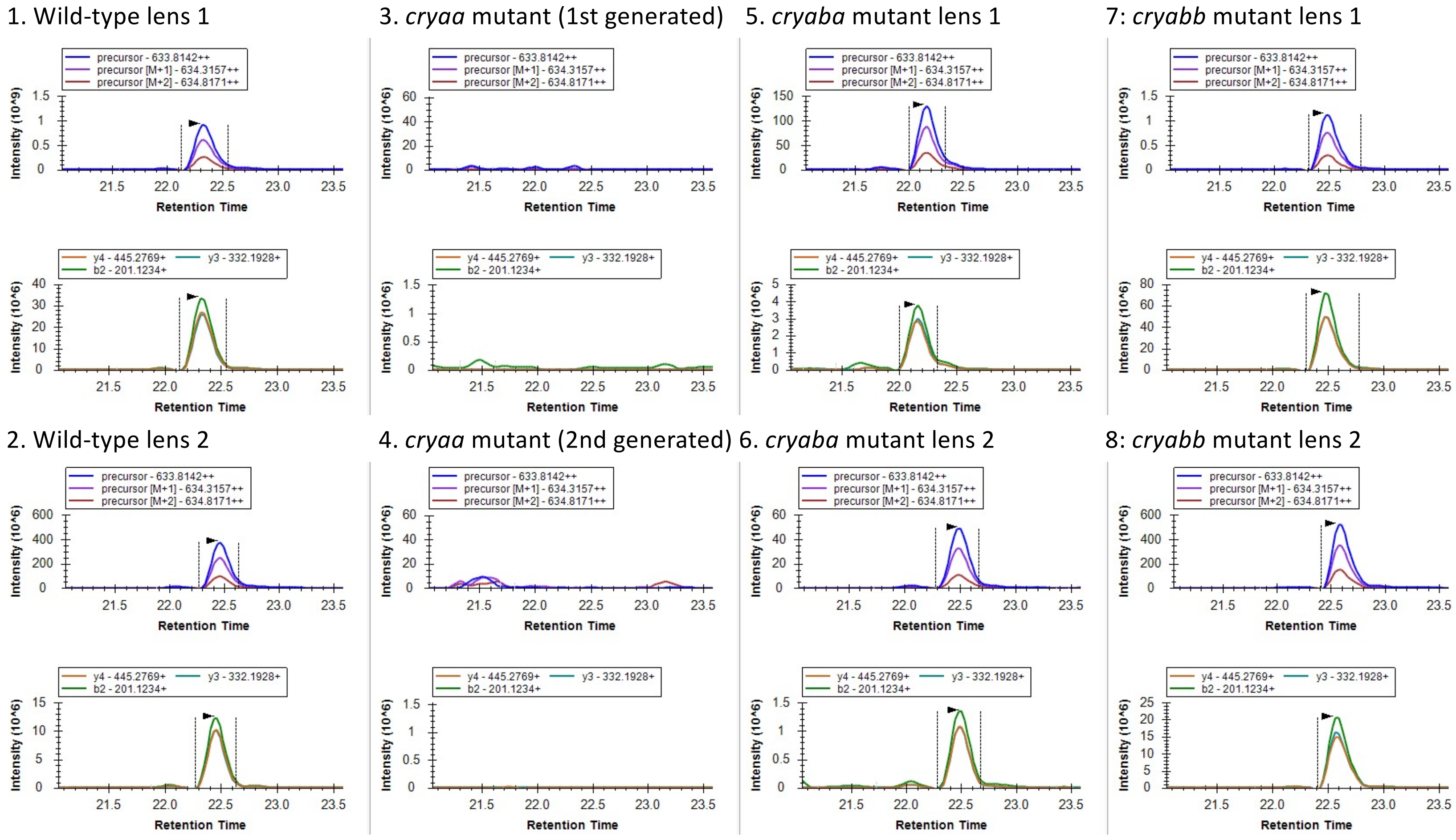

C) αA: R.LPSNVDQSAIT**C**TLSADGLLTL**C**GPK.T [121, 146]

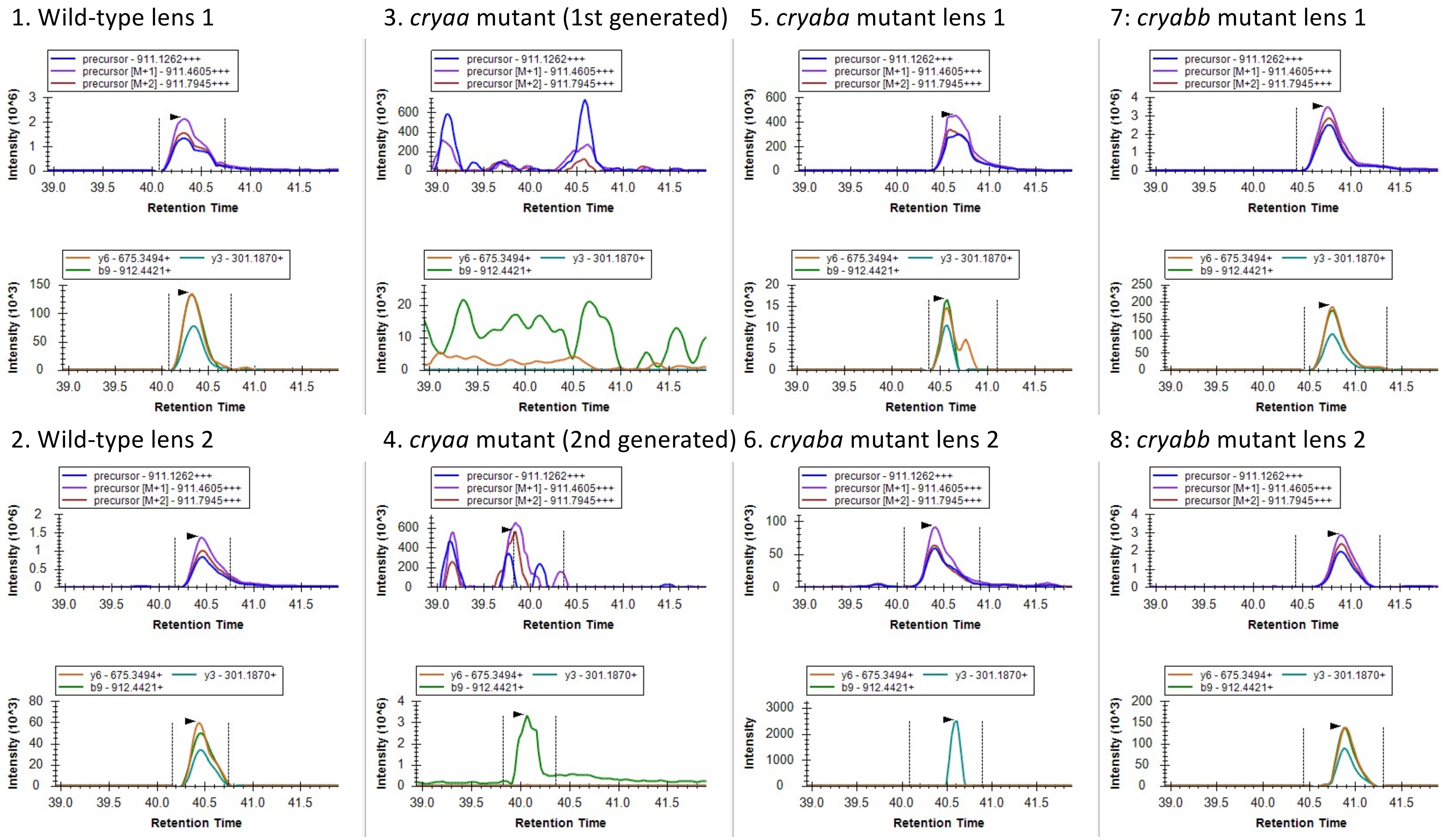

### D) αBa: K.HFSPDELTVK.V [80, 89]

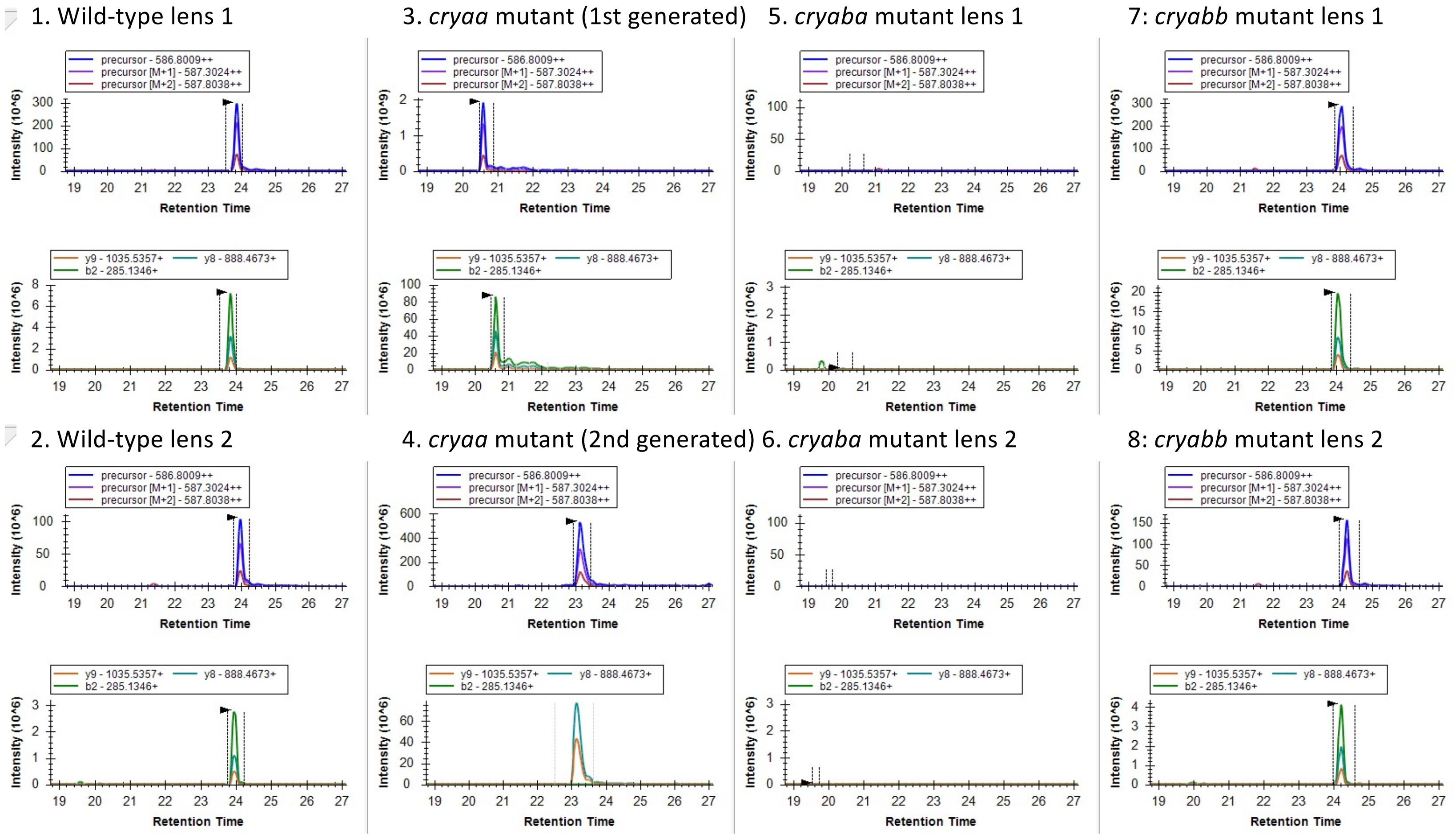

### E) αBa: K.VNEDFIEIHGK.H [90, 100]

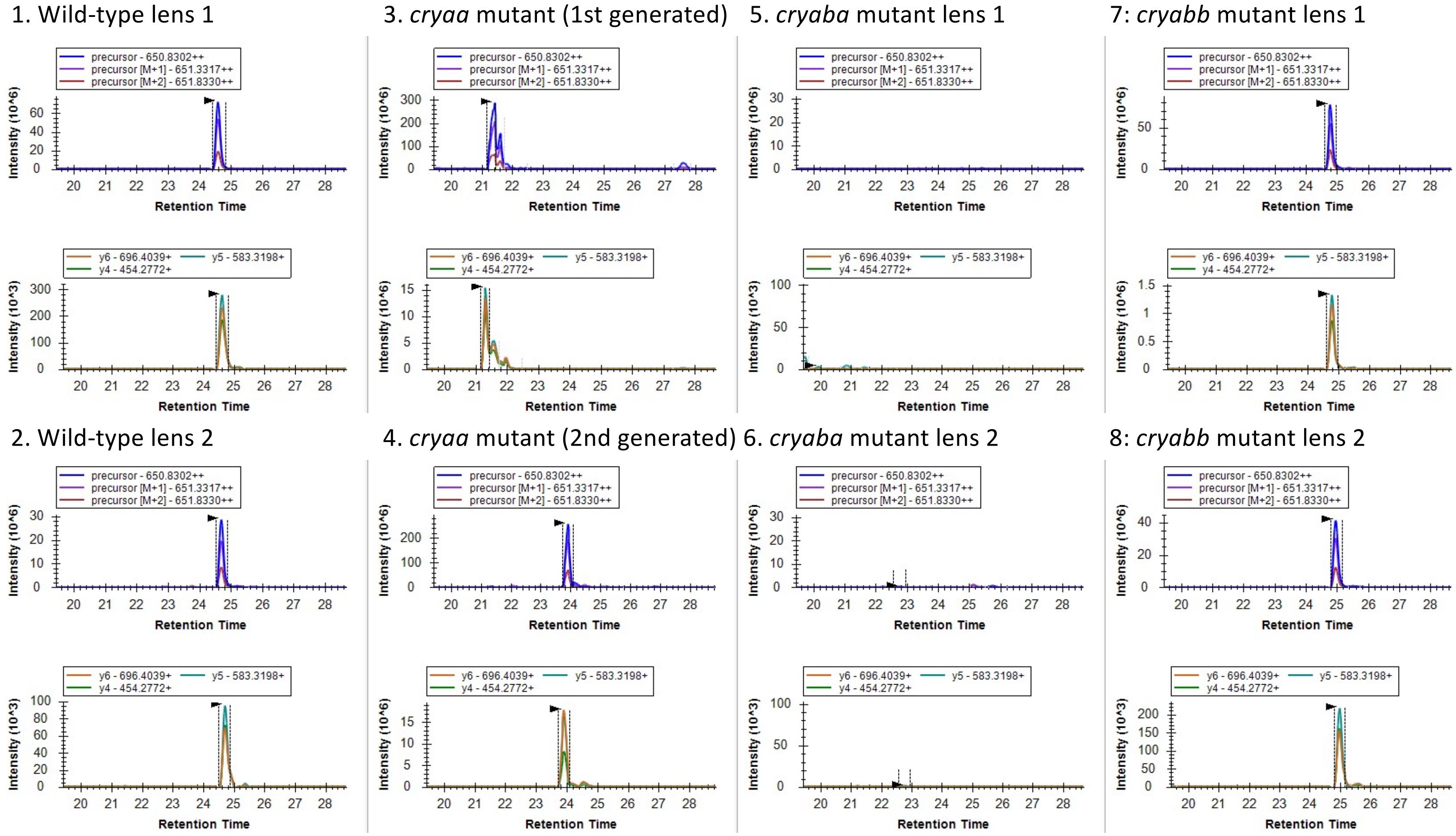

F) αBa: R.SIPII**C**GEKPPAQK.- [155, 168]

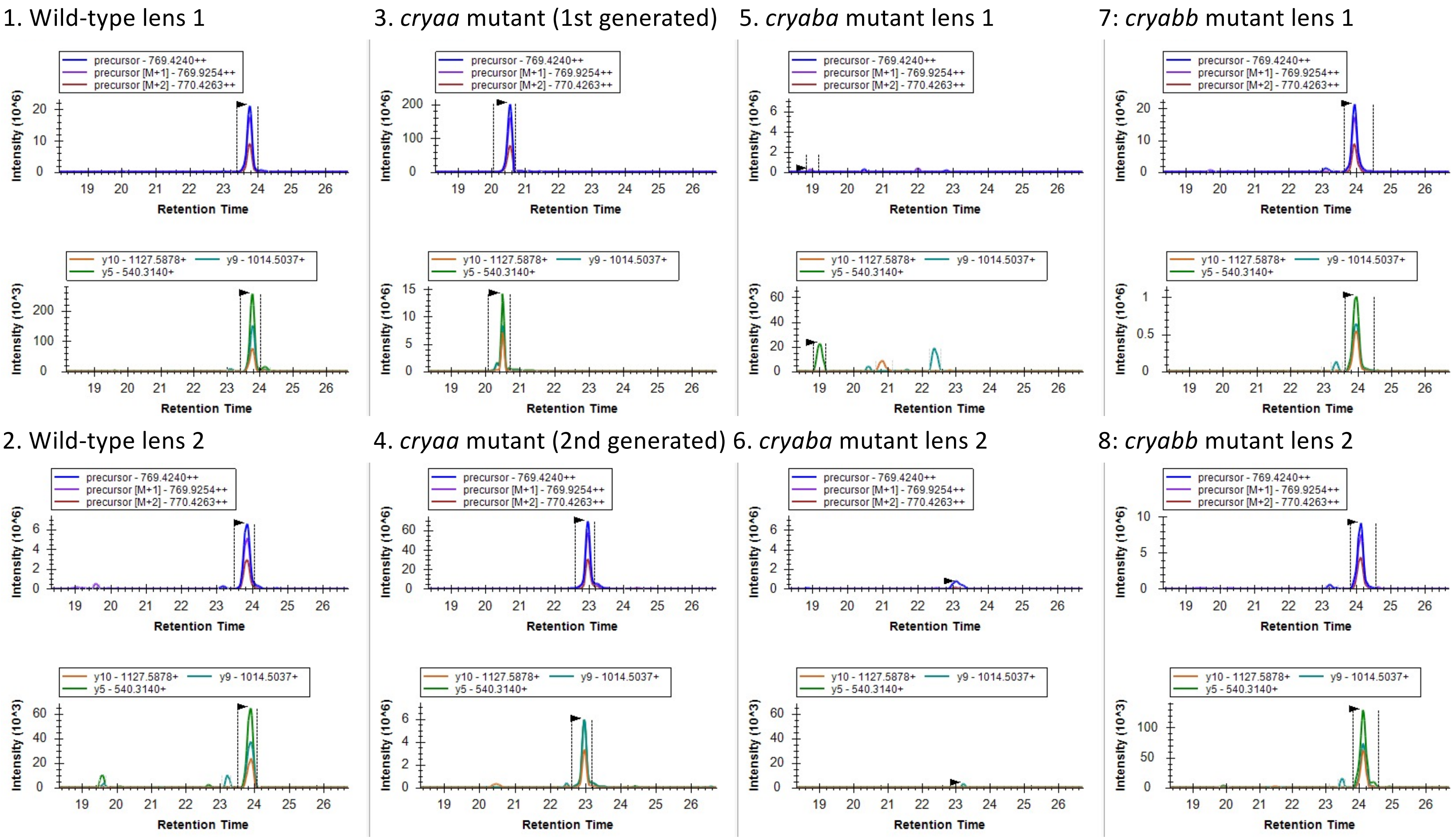

#### G) αBb: R.ILFPIFFPR.R [12, 20]

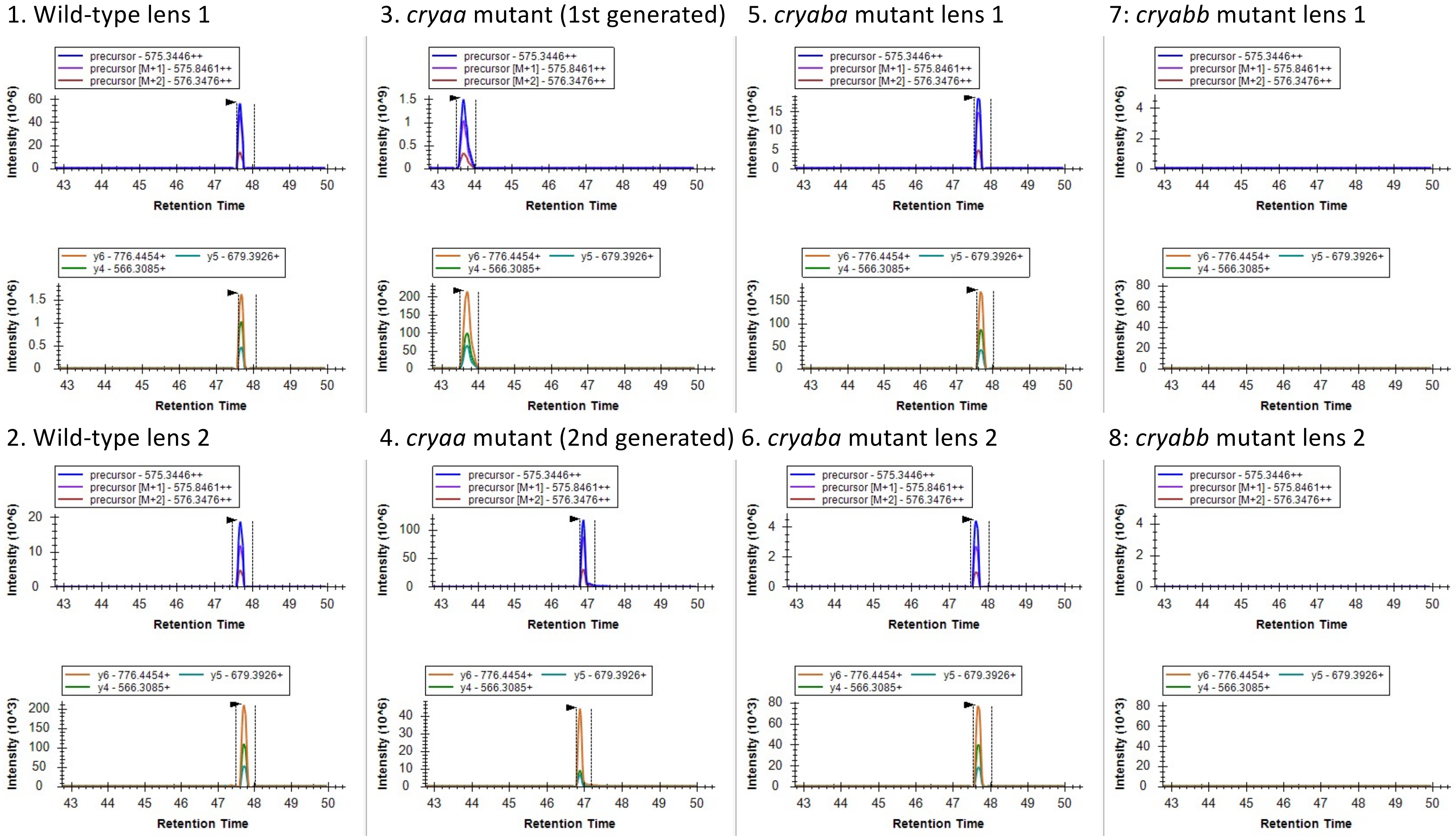

H) αBb: R.SPSWMESGVSEVK.M [46, 58]

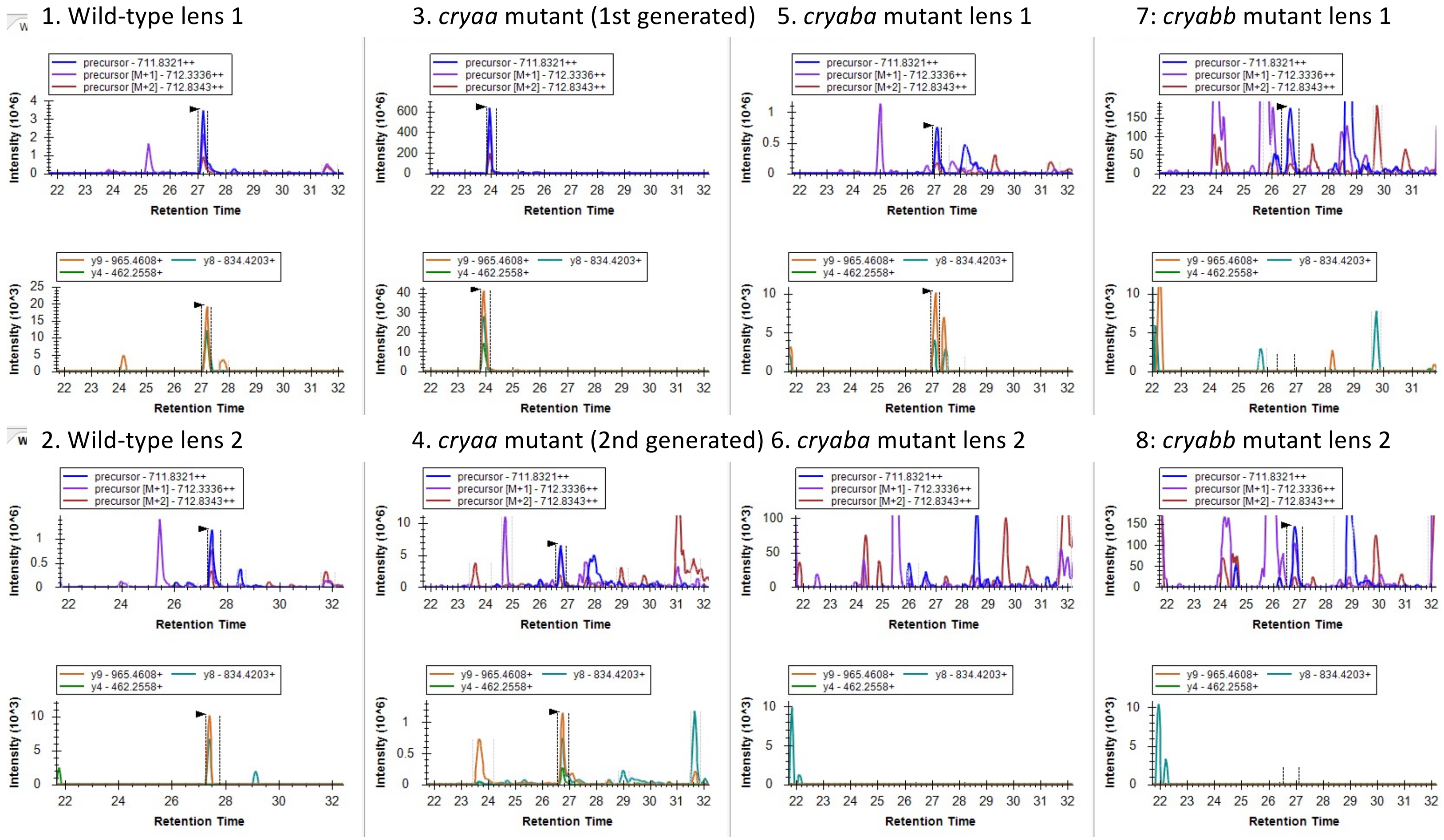

I) αBb: K.IIGDFIEIHAK.H [82, 92]

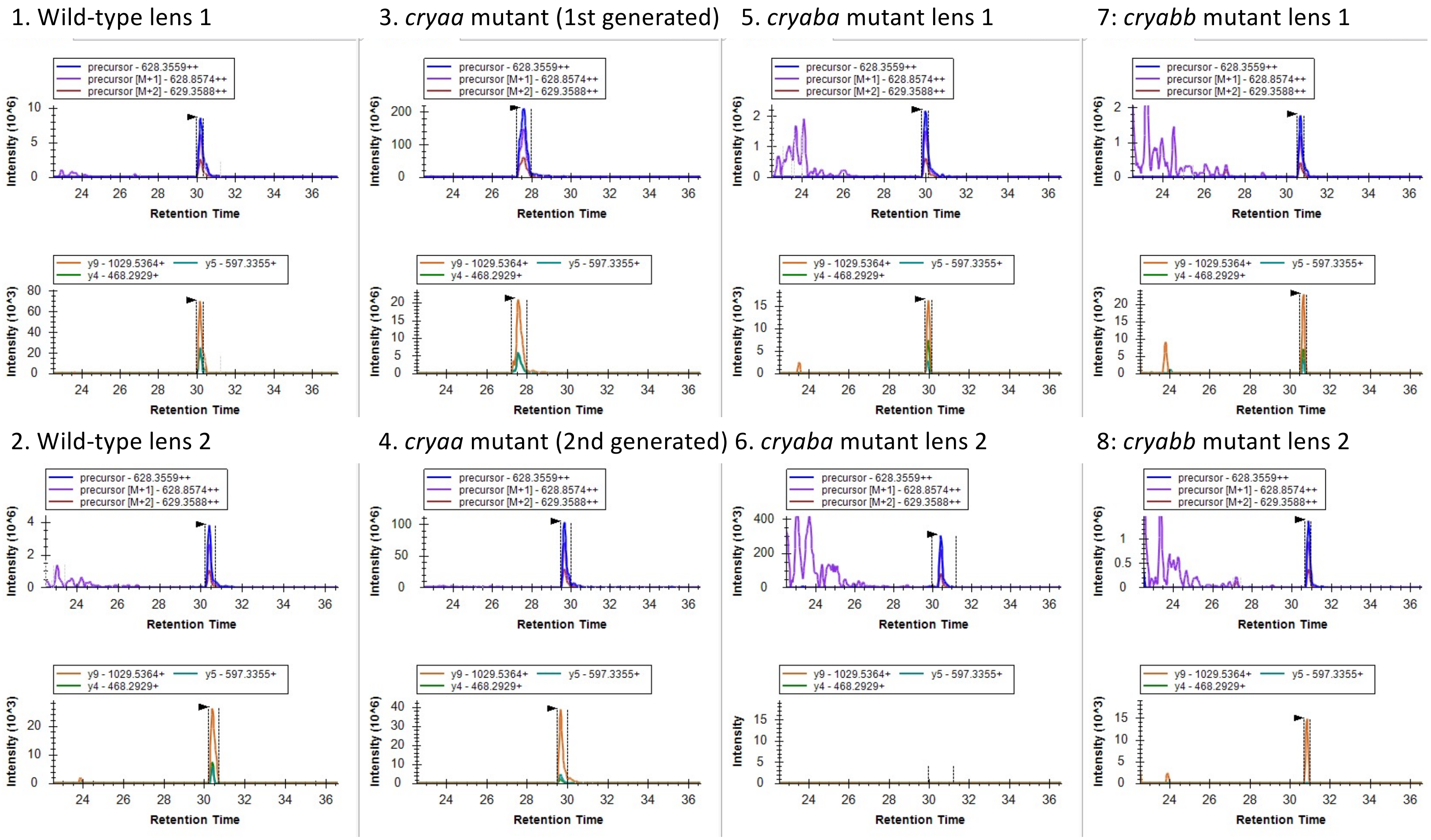
