## Supplemental Figure 2 for "Effects of α-crystallin gene knockout on zebrafish lens development"

### Supplementary Figure 2

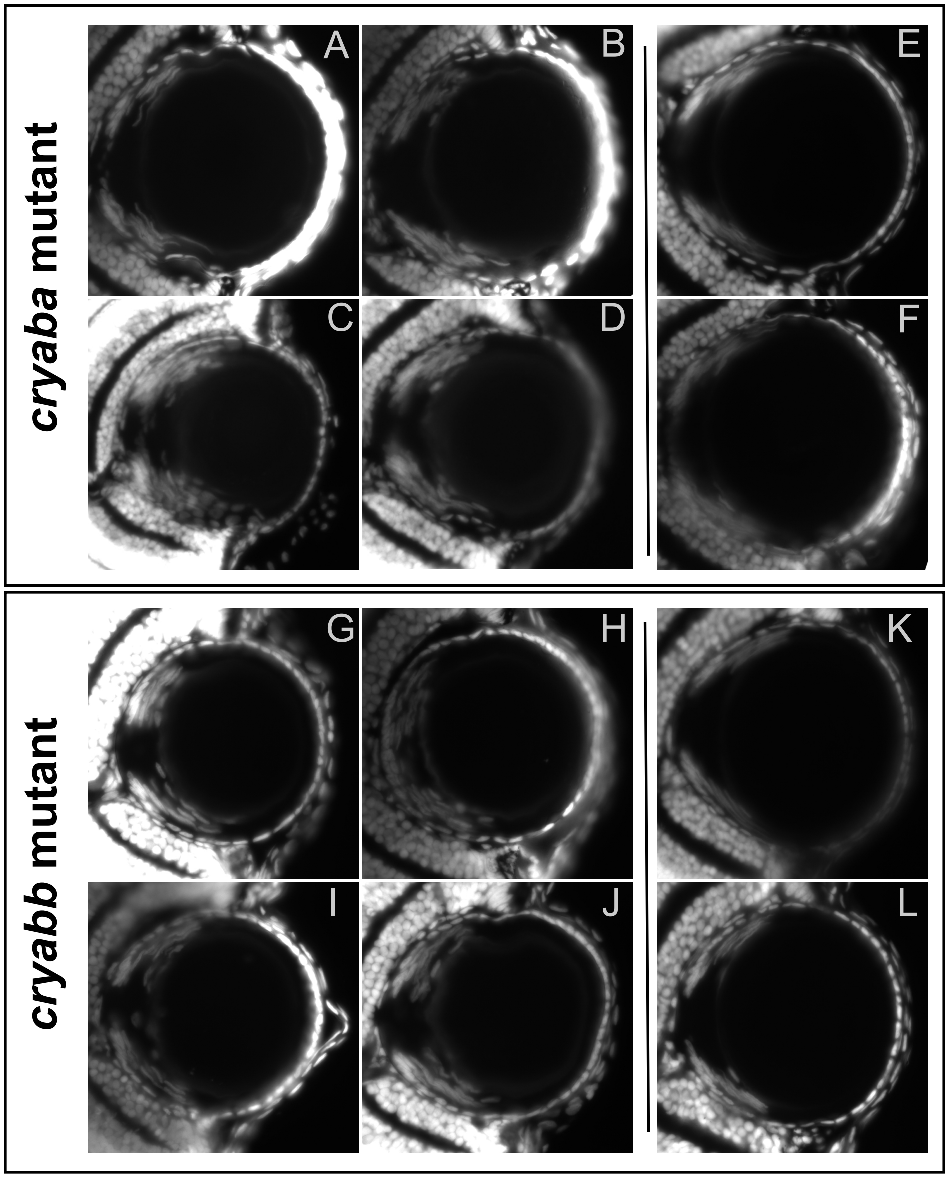


**Supplementary Figure 2.** Cryosectioned lenses at 3 and 4 dpf stained with DAPI to assess clearance of fiber cell nuclei. Four representative lenses are shown for *cryaba* and *cryabb* mutant lines at 3 dpf (A-D and G-J), and two representative lenses at 4 dpf (E-F and K-L). Most images were taken at 200X total magnification, with some taken at 100X to capture more lens cell nuclei in one focal plane. Retina is oriented to the left in each image, with cornea to the right. The four images at 3 dpf for each genotype demonstrate the range of phenotypes found, with the top left panel the most normal and bottom right the most abnormal.
