## Supplemental Figure 3 for "Effects of α-crystallin gene knockout on zebrafish lens development"

### Supplementary Figure 3

A

B

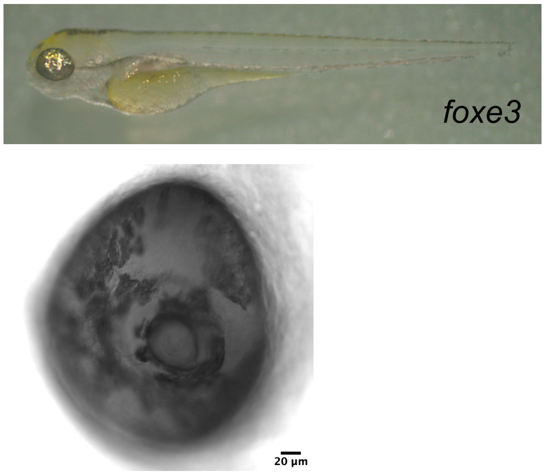


**Supplementary Figure 3.** Effects of injecting a four gRNA mix targeting *foxe3*. Injected crispant fish at 3 dpf showing lack of a pupil characteristic of a previously reported phenotype from *foxe3* null larvae (A; Krall and Lachke 2018). Lens from *foxe3* crispant larva showing severe eye phenotype with DIC optics (B). The narrowing of the pupil makes it difficult to see the edges of the lens.
