## Supplemental Table 1 for "Effects of α-crystallin gene knockout on zebrafish lens development"

**Supplemental Table 1. Primers used to generate gRNAs and amplicons for genotyping.** The NCBI Gene ID is shown for each alpha-crystallin gene. *cryaa* primers are shown for the single gRNA used to generate the original knockout line, the double gRNAs used for the subsequent deletion knockout line, and the four-gRNA mix (poly gRNAs) used to screen for phenotypes directly in the injected embryos. The gene-specific target region within each primer is shown in bold.

| **Primer** | **Primer Sequence** |
| --- | --- |
| Universal Scaffold | AAAAGCACCGACTCGGTGCCACTTTTTCAAGTTGATAACGGACTAGCCTTATTTTAACTTGCTATTTCTAGCTCTAAAAC |
| ***cryaa* Gene ID 100000769** | |
| Single gRNA | TAATACGACTCACTATAGG**GTTCGATTATGACCTATT**GTTTTAGAGCTAGAAATAGCAAG |
| Double gRNA #1 | TAATACGACTCACTATAGG**ATCTTCAGAGAAACTTGCAT**GTTTTAGAGCTAGAAATAGCAAG |
| Double gRNA #2 | TAATACGACTCACTATAGG**TTGGTTCAGACGCACACTGG**GTTTTAGAGCTAGAAATAGCAAG |
| Poly gRNA #1 | TAATACGACTCACTATAGG**TTGGTTCAGACGCACACT**GTTTTAGAGCTAGAAATAGC |
| Poly gRNA #2 | TAATACGACTCACTATAGG**AGTGAGTGTCGATAGTAA**GTTTTAGAGCTAGAAATAGC |
| Poly gRNA #3 | TAATACGACTCACTATAGG**CTTTCTCCATGCTTGCCC**GTTTTAGAGCTAGAAATAGC |
| Poly gRNA #4 | TAATACGACTCACTATAGG**TAGCGACGATGGAACTCA**GTTTTAGAGCTAGAAATAGC |
| ***cryaba* Gene ID 30393** | |
| Single gRNA | TAATACGACTCACTATAGG**TTGTTCCCAGGCTTCTTC**GTTTTAGAGCTAGAAATAGCAAG |
| ***cryabb* Gene ID 436943** | |
| Single gRNA | TAATACGACTCACTATAGG**TTCCGGCGCATCTTATTT**GTTTTAGAGCTAGAAATAGCAAG |
| **PCR Primers for Genotyping (annealing temp shown in parentheses)** | |
| *cryaa* forward | TTCATTCCACTGTGGAGACC (62°C) |
| *cryaa* reverse | CACCTGAGTTGGAGGAGTCC |
| *cryaba* forward | ACTGGAGATAGATCAGTGGCTGAC (55°C) |
| *cryaba* reverse | GGCAGGTTAGGGTAATTAGGCAAG |
| *cryabb* forward | AGAAGATTTGCAGAAGAGGC (55°C) |
| *cryabb* reverse | GAAGAAAGGAAGATCGTTGG |
